# Urinary bladder enlargement across nine rodent models of diabetes: correlations with glucose and insulin levels

**DOI:** 10.1101/2021.10.25.465601

**Authors:** Zeynep E. Yesilyurt, Jan Matthes, Edith Hintermann, Tamara R. Castañeda, Ralf Elvert, Jesus H. Beltran-Ornelas, Diana L. Silva-Velasco, Ning Xia, Aimo Kannt, Urs Christen, David Centurion, Huige Li, Andrea Pautz, Ebru Arioglu-Inan, Martin C. Michel

## Abstract

The urinary bladder is markedly enlarged in the type 1 diabetes mellitus model of streptozotocin (STZ)-injected rats, but much less data exist for models of type 2 diabetes (T2DM). Diabetic polyuria has been proposed to explain bladder enlargement. We have collected data on bladder weight and blood glucose from 16 studies representing 9 distinct rodent diabetes (7 T2DM) and obesity models; some included arms with diets and/or pharmacological treatments. Data were analyzed for bladder enlargement and for correlations between bladder weight on the one and glucose levels on the other hand. Our data confirm major bladder enlargements in STZ rats, minor if any enlargement in fructose-fed rats, db/db mice and mice on a high-fat diet. For the first time we report bladder weight data on 5 other models with presence and degree of bladder enlargement being heterogeneous across models. Bladder weight was correlated with plasma glucose in some but not other models, but correlations were moderate to weak except for RIP-LMCV mice. We conclude that the presence and extent of bladder enlargement varies markedly across diabetes models, particularly TD2M models; our data do not support the idea that bladder enlargement is primarily driven by glucose levels/glucosuria.

## Introduction

Diabetes mellitus is a healthcare challenge with globally increasing prevalence and causes major morbidity and mortality related to cardiovascular, renal and ocular function (1). Dysfunction of the lower urinary tract in general and of the bladder in particular are at least as common, occurring in 80% and 50% of diabetic patients, respectively (2). For example, lower urinary tract dysfunction linked to benign prostatic enlargement increases with age; men with concomitant diabetes have worse symptoms that are equivalent to non-diabetic men who are 12 years older (3). Moreover, men with lower urinary tract symptoms and concomitant diabetes are 37% more likely to experience progression of urinary dysfunction (4). While lower urinary tract dysfunction does not lead to major morbidity or mortality, it reduces the quality of life of the afflicted patients (5, 6) and their partners (7) by impairing social interactions during the day and sleep during the night. Moreover, urinary tract dysfunction is associated with emergency room visits, hospitalizations and loss of work productivity (8).

The pathophysiology of lower urinary tract dysfunction in diabetes is poorly understood and dedicated therapeutic strategies other than normalizing glucose levels are lacking. According to a systematic review of >70 studies, bladder enlargement is consistently found in the streptozotocin (STZ)-induced rat model of type 1 diabetes mellitus (T1DM), by average resulting in a doubling of bladder weight (BW) (9). While studied much less frequently, a comparable enlargement of the urinary bladder appears to exist in all other T1DM models that have been tested (10). Despite type 2 diabetes (T2DM) being considerably more prevalent than T1DM, much fewer studies have explored bladder enlargement in animal models of T2DM and have yielded conflicting results: it was comparable to that in STZ-injected rats in Zucker diabetic fatty rats, present but less pronounced in fructose-fed rats and db/db mice and absent in Goto Kakizaki rats or mice on a high-fat diet (HFD), the latter primarily being an obesity and not necessarily a diabetes model; of note, each of the T2DM/obesity models has been tested in few studies only (10). Thus, it remains unclear whether bladder enlargement occurs in diabetes in general, is restricted to T1DM models or occurs in some but not all T2DM models. Treatment with insulin prevents or, when started in established diabetes, reverses bladder enlargement in STZ-injected rats (9). However, no treatment studies have reported effects on BW in animal models of T2DM or with treatments other than insulin in those of T1DM.

The pathophysiology underlying bladder enlargement in animal models of diabetes is largely unknown. A leading theory is that increased glucose levels act as an osmotic diuretic when exceeding the renal reabsorption threshold of 9-10 mM and that the bladder enlarges as a response to increased urine flow (diabetic polyuria). This hypothesis is supported by studies in rats in which chronic treatment with an osmotic diuretic consistently yielded similar degrees of diuresis and of bladder enlargement as compared to STZ injection, despite not affecting glucose levels (11–17). This theory implies that bladder enlargement is correlated to blood glucose levels if these exceed the renal reabsorption threshold. However, this theory has been questioned because only moderate if any correlation between degree of bladder enlargement and blood glucose levels were found in an inter-study comparison at the group level (10) or based on data from individual animals within one study (18).

Considering this background, our study had three specific aims: The primary aim was to explore presence and extent of bladder enlargement across a wide range of rodent models of diabetes, particularly of T2DM. The second was to explore the effects of diets and pharmacological treatments other than insulin on bladder enlargement. The third aim was to explore correlations between blood glucose levels and bladder weight in various models based on intra-study comparisons at the individual animal level. To address these aims, we have collected data on glucose levels (in some cases also insulin levels), bladder and body weight from various ongoing studies primarily designed to address questions unrelated to the urinary bladder. This has allowed us to use data from 16 studies with 2-8 arms each representing 9 distinct rat and mouse models and a total of 513 animals without sacrificing a single animal for the purpose of our study. Taken together we present what may be the most comprehensive inter-model comparison ever reported for any parameter in diabetes. This highlights that major *in vivo* scientific advancement can be made while markedly reducing animal use.

## Results

### Model characterization

#### Glycemic state

Based on our operational definition of normoglycemia (blood glucose concentrations <8 mM), hyperglycemia (8-16 mM), and overt diabetes (>16 mM), the control groups in both STZ rat studies, in both ZSF1 rat studies, in all three fructose-fed rat studies, and in the rats with neonatal STZ injection were normoglycemic; in contrast, control groups of the rat insulin promotor lymphocytic choriomeningitis virus (RIP-LMCV mice) were hyperglycemic; control C57BL/6J were normoglycemic in two and hyperglycemic in two other studies, and C57BL/6N mice normoglycemic in one and hyperglycemic in another study (Table 1). STZ rats, RIP-LMCV mice, obese 28-weeks old ZSF1 rats, insulin receptor substrate 2 (IRS2) knock-out mice and db/db mice were diabetic; 20-weeks old ZSF1 rats, rats with neonatal STZ injection, ob/ob mice and HFD mice were hyperglycemic; fructose-fed rats had somewhat greater glucose levels than their control in one, but not in two other studies and did not reach the above threshold of hyperglycemia in any study (Table 1).

**Table 1:**
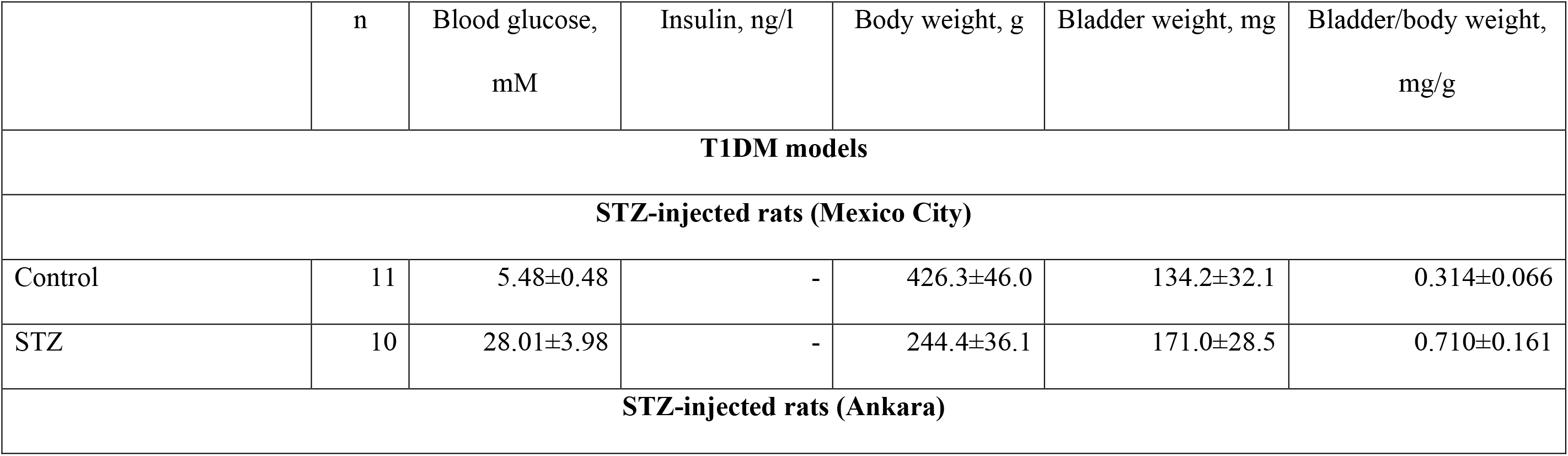
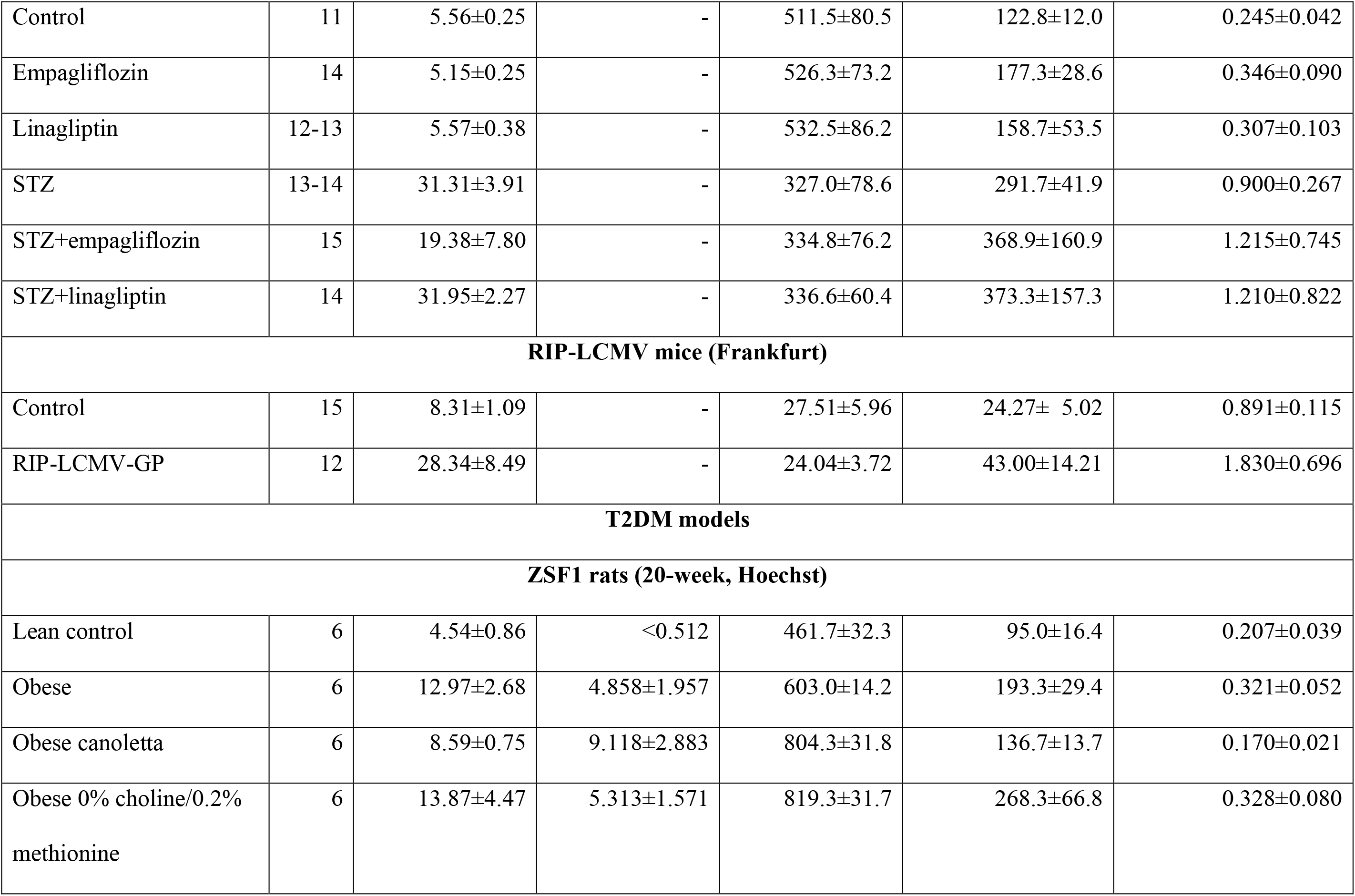
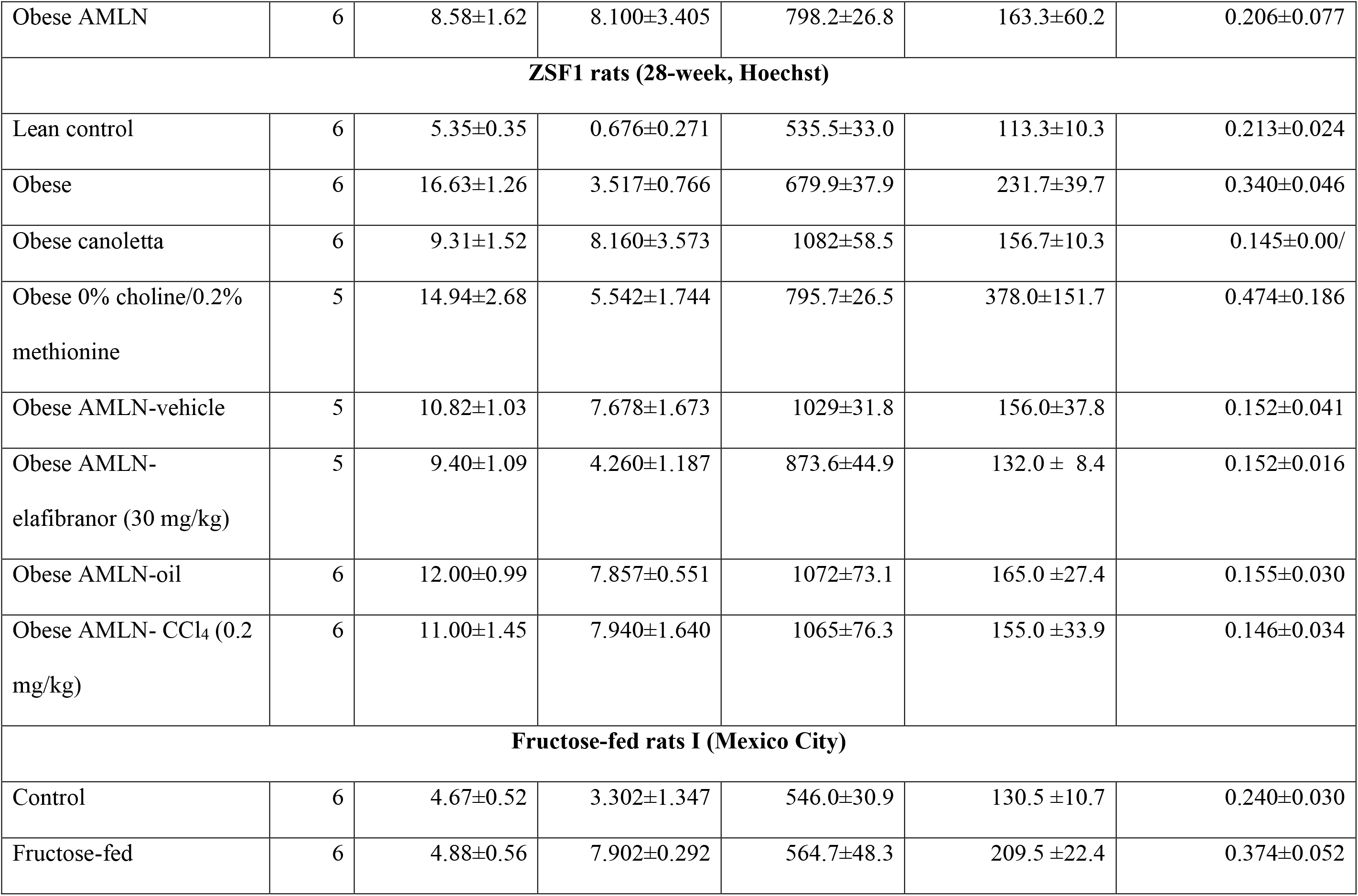
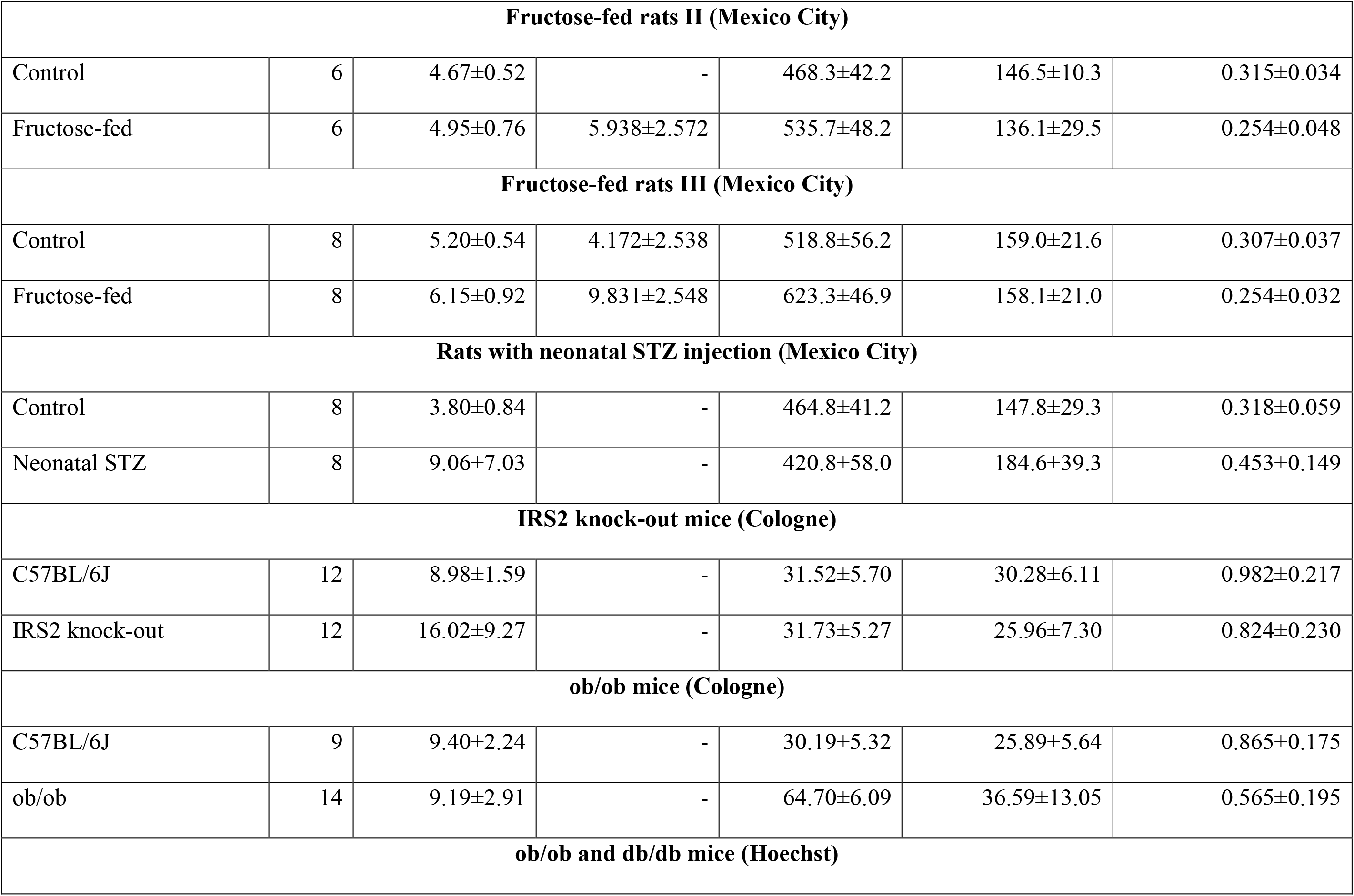
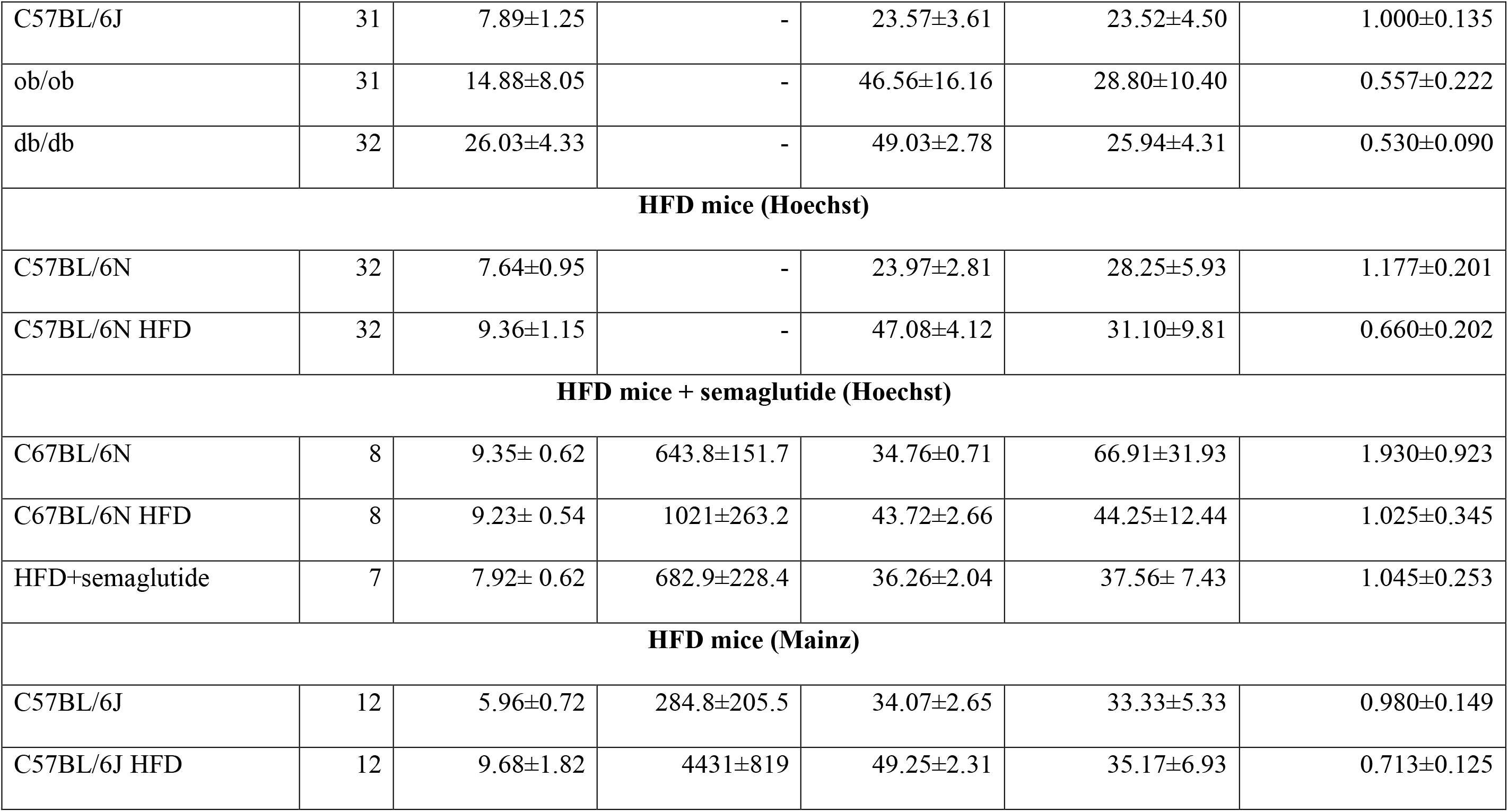
Blood glucose, insulin (selected studies only), body weight, bladder weight, and bladder/body weight across animal models. Data are shown as means ± SD of the indicated number of animals. Insulin concentrations were below detection limit (0.000512 ng/ml) in lean ZSF1 rats in all animals in the 20- and 4/6 in the 28-week study; for calculation purposes they were set to 0.000512 ng/ml. Data from each individual animal of each study are shown in the Online Supplement.

|  | n | Blood glucose,<br>mM | Insulin, ng/l | Body weight, g | Bladder weight, mg | Bladder/body weight,<br>mg/g |
| --- | --- | --- | --- | --- | --- | --- |
| **T1DM models** |  |  |  |  |  |  |
| **STZ-injected rats (Mexico City)** |  |  |  |  |  |  |
| Control | 11 | 5.48±0.48 | - | 426.3±46.0 | 134.2±32.1 | 0.314±0.066 |
| STZ | 10 | 28.01±3.98 | - | 244.4±36.1 | 171.0±28.5 | 0.710±0.161 |
| **STZ-injected rats (Ankara)** |  |  |  |  |  |  |
| Control | 11 | 5.56±0.25 | - | 511.5±80.5 | 122.8±12.0 | 0.245±0.042 |
| Empagliflozin | 14 | 5.15±0.25 | - | 526.3±73.2 | 177.3±28.6 | 0.346±0.090 |
| Linagliptin | 12-13 | 5.57±0.38 | - | 532.5±86.2 | 158.7±53.5 | 0.307±0.103 |
| STZ | 13-14 | 31.31±3.91 | - | 327.0±78.6 | 291.7±41.9 | 0.900±0.267 |
| STZ+empagliflozin | 15 | 19.38±7.80 | - | 334.8±76.2 | 368.9±160.9 | 1.215±0.745 |
| STZ+linagliptin | 14 | 31.95±2.27 | - | 336.6±60.4 | 373.3±157.3 | 1.210±0.822 |
| **RIP-LCMV mice (Frankfurt)** |  |  |  |  |  |  |
| Control | 15 | 8.31±1.09 | - | 27.51±5.96 | 24.27± 5.02 | 0.891±0.115 |
| RIP-LCMV-GP | 12 | 28.34±8.49 | - | 24.04±3.72 | 43.00±14.21 | 1.830±0.696 |
| **T2DM models** |  |  |  |  |  |  |
| **ZSF1 rats (20-week, Hoechst)** |  |  |  |  |  |  |
| Lean control | 6 | 4.54±0.86 | <0.512 | 461.7±32.3 | 95.0±16.4 | 0.207±0.039 |
| Obese | 6 | 12.97±2.68 | 4.858±1.957 | 603.0±14.2 | 193.3±29.4 | 0.321±0.052 |
| Obese canoletta | 6 | 8.59±0.75 | 9.118±2.883 | 804.3±31.8 | 136.7±13.7 | 0.170±0.021 |
| Obese 0% choline/0.2% methionine | 6 | 13.87±4.47 | 5.313±1.571 | 819.3±31.7 | 268.3±66.8 | 0.328±0.080 |
| Obese AMLN | 6 | 8.58±1.62 | 8.100±3.405 | 798.2±26.8 | 163.3±60.2 | 0.206±0.077 |
| **ZSF1 rats (28-week, Hoechst)** |  |  |  |  |  |  |
| Lean control | 6 | 5.35±0.35 | 0.676±0.271 | 535.5±33.0 | 113.3±10.3 | 0.213±0.024 |
| Obese | 6 | 16.63±1.26 | 3.517±0.766 | 679.9±37.9 | 231.7±39.7 | 0.340±0.046 |
| Obese canoletta | 6 | 9.31±1.52 | 8.160±3.573 | 1082±58.5 | 156.7±10.3 | 0.145±0.00/ |
| Obese 0% choline/0.2% methionine | 5 | 14.94±2.68 | 5.542±1.744 | 795.7±26.5 | 378.0±151.7 | 0.474±0.186 |
| Obese AMLN-vehicle | 5 | 10.82±1.03 | 7.678±1.673 | 1029±31.8 | 156.0±37.8 | 0.152±0.041 |
| Obese AMLN-elafibranor (30 mg/kg) | 5 | 9.40±1.09 | 4.260±1.187 | 873.6±44.9 | 132.0 ± 8.4 | 0.152±0.016 |
| Obese AMLN-oil | 6 | 12.00±0.99 | 7.857±0.551 | 1072±73.1 | 165.0 ±27.4 | 0.155±0.030 |
| Obese AMLN- CCl<sub>4</sub> (0.2 mg/kg) | 6 | 11.00±1.45 | 7.940±1.640 | 1065±76.3 | 155.0 ±33.9 | 0.146±0.034 |
| **Fructose-fed rats I (Mexico City)** |  |  |  |  |  |  |
| Control | 6 | 4.67±0.52 | 3.302±1.347 | 546.0±30.9 | 130.5 ±10.7 | 0.240±0.030 |
| Fructose-fed | 6 | 4.88±0.56 | 7.902±0.292 | 564.7±48.3 | 209.5 ±22.4 | 0.374±0.052 |
| **Fructose-fed rats II (Mexico City)** |  |  |  |  |  |  |
| Control | 6 | 4.67±0.52 | - | 468.3±42.2 | 146.5±10.3 | 0.315±0.034 |
| Fructose-fed | 6 | 4.95±0.76 | 5.938±2.572 | 535.7±48.2 | 136.1±29.5 | 0.254±0.048 |
| **Fructose-fed rats III (Mexico City)** |  |  |  |  |  |  |
| Control | 8 | 5.20±0.54 | 4.172±2.538 | 518.8±56.2 | 159.0±21.6 | 0.307±0.037 |
| Fructose-fed | 8 | 6.15±0.92 | 9.831±2.548 | 623.3±46.9 | 158.1±21.0 | 0.254±0.032 |
| **Rats with neonatal STZ injection (Mexico City)** |  |  |  |  |  |  |
| Control | 8 | 3.80±0.84 | - | 464.8±41.2 | 147.8±29.3 | 0.318±0.059 |
| Neonatal STZ | 8 | 9.06±7.03 | - | 420.8±58.0 | 184.6±39.3 | 0.453±0.149 |
| **IRS2 knock-out mice (Cologne)** |  |  |  |  |  |  |
| C57BL/6J | 12 | 8.98±1.59 | - | 31.52±5.70 | 30.28±6.11 | 0.982±0.217 |
| IRS2 knock-out | 12 | 16.02±9.27 | - | 31.73±5.27 | 25.96±7.30 | 0.824±0.230 |
| **ob/ob mice (Cologne)** |  |  |  |  |  |  |
| C57BL/6J | 9 | 9.40±2.24 | - | 30.19±5.32 | 25.89±5.64 | 0.865±0.175 |
| ob/ob | 14 | 9.19±2.91 | - | 64.70±6.09 | 36.59±13.05 | 0.565±0.195 |
| **ob/ob and db/db mice (Hoechst)** |  |  |  |  |  |  |
| C57BL/6J | 31 | 7.89±1.25 | - | 23.57±3.61 | 23.52±4.50 | 1.000±0.135 |
| ob/ob | 31 | 14.88±8.05 | - | 46.56±16.16 | 28.80±10.40 | 0.557±0.222 |
| db/db | 32 | 26.03±4.33 | - | 49.03±2.78 | 25.94±4.31 | 0.530±0.090 |
| **HFD mice (Hoechst)** |  |  |  |  |  |  |
| C57BL/6N | 32 | 7.64±0.95 | - | 23.97±2.81 | 28.25±5.93 | 1.177±0.201 |
| C57BL/6N HFD | 32 | 9.36±1.15 | - | 47.08±4.12 | 31.10±9.81 | 0.660±0.202 |
| **HFD mice + semaglutide (Hoechst)** |  |  |  |  |  |  |
| C67BL/6N | 8 | 9.35± 0.62 | 643.8±151.7 | 34.76±0.71 | 66.91±31.93 | 1.930±0.923 |
| C67BL/6N HFD | 8 | 9.23± 0.54 | 1021±263.2 | 43.72±2.66 | 44.25±12.44 | 1.025±0.345 |
| HFD+semaglutide | 7 | 7.92± 0.62 | 682.9±228.4 | 36.26±2.04 | 37.56± 7.43 | 1.045±0.253 |
| **HFD mice (Mainz)** |  |  |  |  |  |  |
| C57BL/6J | 12 | 5.96±0.72 | 284.8±205.5 | 34.07±2.65 | 33.33±5.33 | 0.980±0.149 |
| C57BL/6J HFD | 12 | 9.68±1.82 | 4431±819 | 49.25±2.31 | 35.17±6.93 | 0.713±0.125 |

Among studies in mice on an HFD, two had increased glucose levels (by about 1.8 and 3.6 mM), but the third did not (Table 1). Multiple diets were tested in obese ZSF1 rats: canoletta lowered glucose levels in 20- and 28-week-old ZSF1 rats by about 4.4 and 7.3 mM, respectively; 0% choline/0.2% methionine diet had no major effects on glucose in either study; Amylin liver NASH (AMLN) diet lowered glucose levels in both studies, although less so when co-administered with oil (vehicle for CCl_4_) (Table 1).

The oral anti-diabetic drug empagliflozin, but not linagliptin, substantially lowered the blood glucose concentration in STZ rats (from 30.31 to 19.38 mM; Table 1) but animals remained diabetic. The PPAR-α/δ agonist elafibranor had no major effect on glucose levels in 28-weeks old ZSF1 rats on an AMLN diet (Table 1). The glucagon-like peptide 1 receptor agonist semaglutide lowered glucose levels in HFD mice by about 1.3 mM, but levels remained in the hyperglycemic range (Table 1).

In all six studies with available insulin data, hyperinsulinemia relative to the respective control were observed (both ZSF1 rat studies, both fructose-fed rat studies, both HFD studies; Table 1). Among treatments, canoletta and AMNL diets further increased insulin concentration in both ZSF1 rat studies, whereas 0% choline/0.2% methionine and elafibranor had no major effect; semaglutide lowered insulin concentration in HFD mice (Table 1).

#### Body weight

Body weight was markedly reduced in STZ rats (>40%) and by <15% in RIP-LCMV mice (Table 1). Among T2DM/obesity models, body weight was markedly increased in ZSF1 rats of either age, in both studies with ob/ob mice, in db/db mice, and in HFD mice (Table 1). Fructose-feeding markedly increased body weight in two studies, but much less so in a third one (Table 1). Rats with neonatal STZ injection and IRS2 knock-out mice did not exhibit major alterations of body weight (Table 1). Empagliflozin and linagliptin had no major effects on body weight, whereas semaglutide normalized body weight and elafibranor reduced it by almost 40% relative to its control (AMNL vehicle; Table 1).

### Bladder enlargement

BW was increased in all T1DM models (both studies with STZ rats and with RIP-LCMV mice) and in some T2DM/obesity models (both studies with ZSF1 rats, one of the three studies with fructose-fed rats, study with rats with neonatal STZ injection, both studies with ob/ob mice and in db/db mice; Table 1, Figure 1). In contrast, no bladder enlargement was observed in the other T2DM models (two out of three studies with fructose-fed rats, IRS2 knock-out mice, all three studies with HFD in mice).

**Figure 1:**
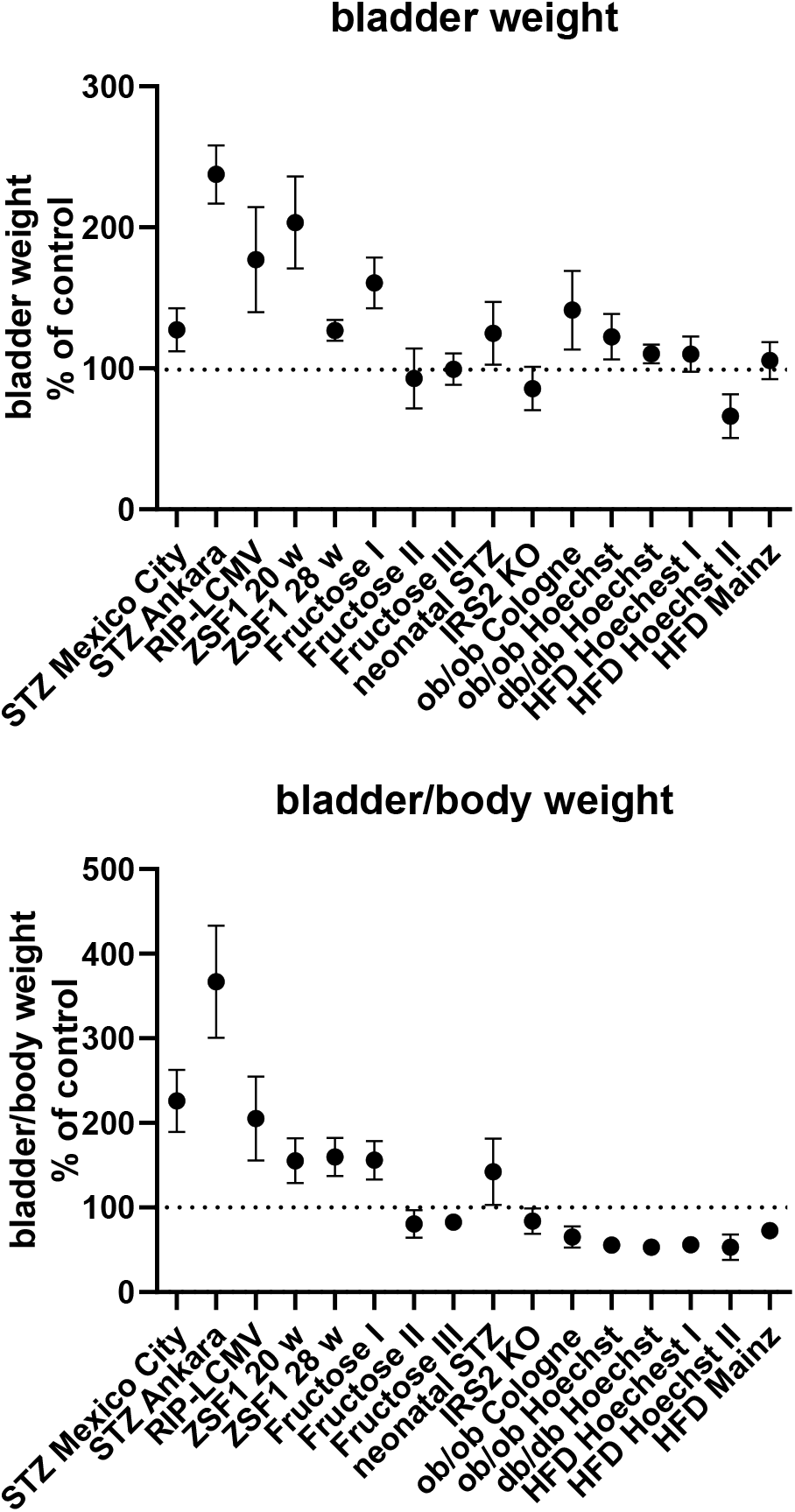
Bladder and bladder/body weight differences across studies. Data are shown as effect sizes comparing the primary hyperglycemic/diabetic vs. the control group expressed as mean difference with its 95% confidence interval. Note that the same control group was used in the calculation of the ob/ob and db/db Hoechst groups.

While HFD did not affect BW in mice (see above), addition of canoletta reduced BW in obese ZSF1 rats assessed at an age of 20 weeks (mean difference -56.7 mg [-86.2; -27.1]), a diet containing 0% choline/0.2% methionine increased BW (mean difference 75.0 mg [CI 8.6; 141.4]), and the AMLN diet had no detectable effect (mean difference -30 mg [CI -91.0; 31.0]; Table 1); however, all three estimates had wide CI making interpretation difficult. Similar effects of the three diets were seen at an age of 28 weeks.

Among pharmacological treatments, empagliflozin and linagliptin led to numerically large increases of BW in STZ rats, but these could not easily be interpreted due to large CI (mean difference 77.2 mg [CI -17.4; 171.8] and 81.6 mg [CI -11.3; 174.5], respectively; Table 1). Elafibranor induced a moderate reduction in BW as compared to obese ZSF1 rats on AMLN diet (mean difference -33 mg [CI -62.0; -4.0]). Semaglutide had no clear effect on BW (mean difference -6.7 mg [CI -18.4; 5.0]).

As body weight exhibited major changes in some of the models (Table 1), a different picture was obtained for bladder/body weight (BBW): An increase was observed in all T1DM models, in both studies with ZSF1 rats, in one of three studies with fructose-fed rats, and in rats with neonatal STZ injection (Table 1, Figure 1). In contrast, reductions of BBW were observed in two of the three studies of fructose-fed rats, both studies in ob/ob mice, in db/db mice and in all three studies with HFD mice; this may primarily be a consequence of increased body weight. No change of BBW was seen in IRS2 knock-out mice.

### Correlations between blood glucose and insulin concentrations and bladder enlargement

Among models with blood glucose levels greater than the renal reabsorption threshold (>16 mM), IRS2 knock-out mice lack and db/db exhibited only a minor increase in BW (Table 1, Figure 1). In contrast, bladder enlargement was observed in one study with a glucose level below the threshold (<9 mM; fructose-fed rat I), whereas studies with glucose levels approximately in the range of the threshold (9-10 mM) exhibited bladder enlargement in two but not in three other studies (Table 1, Figure 1). These findings casted doubt on the diabetic polyuria hypothesis.

To further explore a role of elevated glucose, acting as an osmotic diuretic, as the cause of changes of BW and BBW, we have performed correlation analysis within each model based on individual animal data (Table 2 and Figure 2; additional correlation plots from each model are shown in the Online Supplement). Strength of correlation between glucose level and BW (expressed as r^2^) varied markedly between models and ranged from 0.7226 in RIP-LCMV mice to 0.005 in one of the HFD mice studies. Except for the two ZSF1 rat studies, all groups had r^2^ values of <0.2, indicating that inter-animal variability of glucose levels, serving as a proxy of diabetic polyuria, statistically accounted for less than 20% of differences in BW. Comparable strength of correlation was found when glucose levels were compared to BBW; however, as a notable exception an r^2^ of 0.674 was found for db/db mice, a model in which BW was not markedly changed but body weight about doubled (Table 2). When data from the hyperglycemic/diabetic animals of all studies were pooled, r^2^ was 0.0621 (Figure 2), indicating that glucose did not explain bladder weight variability in an inter-model analysis.

**Table 2:** Correlation between blood glucose and bladder and bladder/body weight across animal models. Animals from diabetic and non-diabetic group were pooled for each correlation analysis. Shown are total number of animals per model, squared correlation coefficient (r^2^) and descriptive p-value. *: negative slope (inverse correlation). A graphical representation of representative groups is shown in Figure 2, all other groups in the Online Supplement.

| n total | Bladder weight |  | Bladder/body weight |  |
| --- | --- | --- | --- | --- |
| | $r^2$ | p | $r^2$ | p |
| **T1DM models** |  |  |  |  |
| **STZ-injected rats (Mexico City)** |  |  |  |  |
| 21 | 0.2346 | 0.0261 | 0.6368 | <0.0001 |
| **STZ-injected rats (Ankara)** |  |  |  |  |
| 79 | 0.3795 | <0.0001 | 0.3220 | <0.0001 |
| **RIP-LCMV mice (Frankfurt)** |  |  |  |  |
| 27 | 0.7226 | <0.0001 | 0.7322 | <0.0001 |
| **T2DM models** |  |  |  |  |
| **ZSF1 rats (20-week, Hoechst)** |  |  |  |  |
| 30 | 0.3632 | 0.0004 | 0.2428 | 0.0057 |
| **ZSF1 rats (28-week, Hoechst)** |  |  |  |  |
| 45 | 0.4127 | <0.0001 | 0.3168 | <0.0001 |
| **Fructose-fed rats I (Mexico City)** |  |  |  |  |
| 12 | 0.0109 | 0.7465 | 0.0044 | 0.8384 |
| **Fructose-fed rats II (Mexico City)** |  |  |  |  |
| 12 | 0.1979* | 0.1473 | 0.2545* | 0.0944 |
| **Fructose-fed rats III (Mexico City)** |  |  |  |  |
| 14 | 0.0488 | 0.4481 | 0.0465* | 0.4590 |
| **Rats with neonatal STZ injection (Mexico City)** |  |  |  |  |
| 16 | 0.3302 | 0.0199 | 0.6262 | 0.0003 |
| **IRS2 knock-out mice (Cologne)** |  |  |  |  |
| 24 | 0.1256 | 0.0893 | 0.1009 | 0.1305 |
| **ob/ob mice (Cologne)** |  |  |  |  |
| 23 | 0.0053 | 0.7410 | 0.0001 | 0.9593 |
| **ob/ob mice (Hoechst)** |  |  |  |  |
| 62 | 0.0339 | 0.1519 | 0.0761* | 0.0300 |
| **db/db mice (Hoechst)** |  |  |  |  |
| 63 | 0.1203 | 0.0054 | 0.6743* | <0.0001 |
| **HFD mice (Hoechst)** |  |  |  |  |
| 64 | 0.0054 | 0.5655 | 0.0383 | 0.1214 |
| **HFD mice + semaglutide (Hoechst)** |  |  |  |  |
| 23 | 0.0787 | 0.1947 | 0.0522 | 0.2945 |
| **HFD mice (Mainz)** |  |  |  |  |
| 24 | 0.0231 | 0.4783 | 0.3614* | 0.0019 |

**Figure 2:**
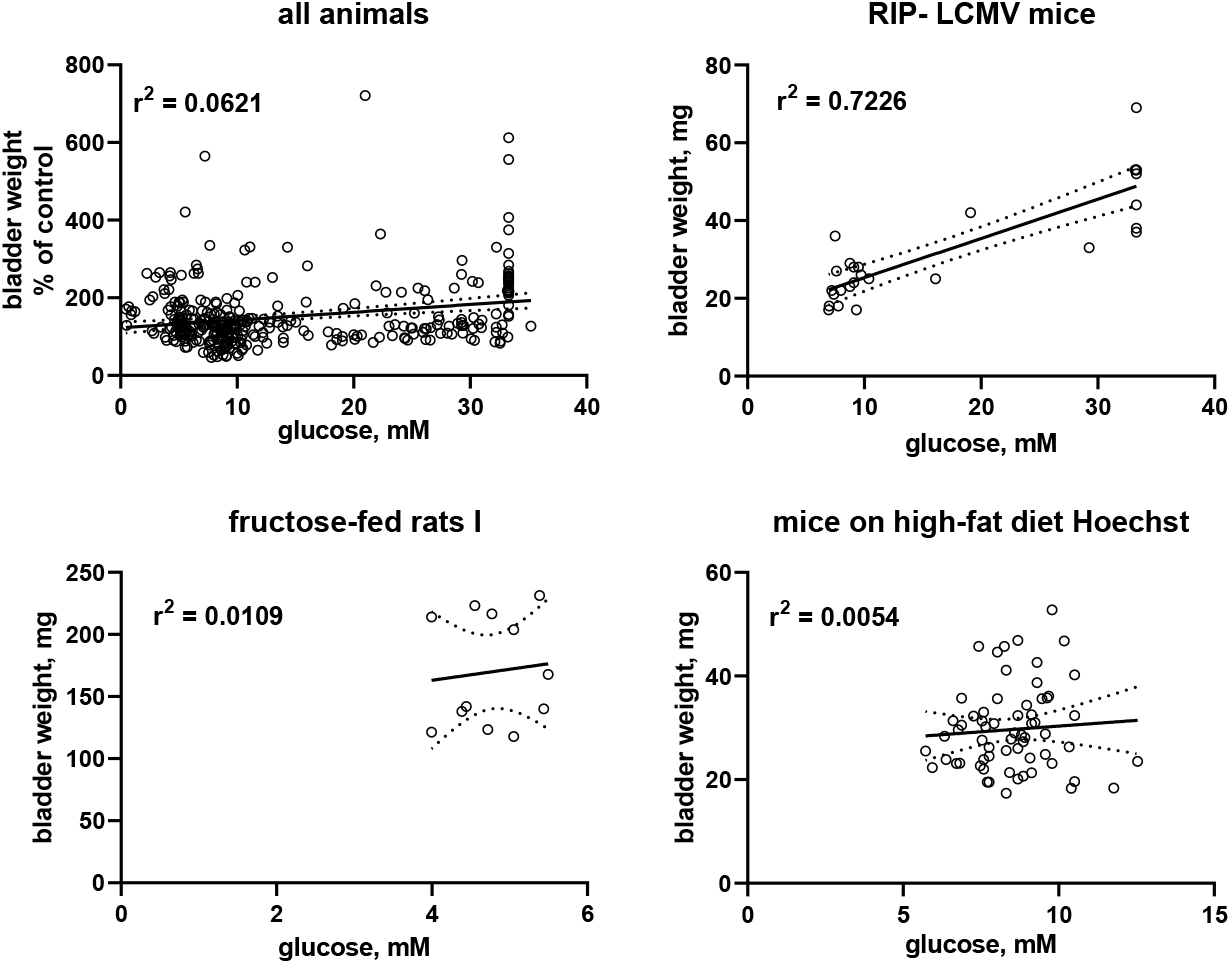
Correlation of bladder and bladder/body weight with glucose levels. To enable pooling of data from all studies, those for the upper left panel shows bladder weight only from the non-control groups expressed as % of mean values in the control group within a study. The other three panels show correlations within three representative studies; data from the remaining studies are shown in the Online Supplement. A quantitative description of the correlations is shown in Table 2. Mean values of bladder weight and glucose level in each study are shown in Table 1.

In addition to the pre-planned correlation analyses for glucose vs. BW, we performed post-hoc correlation analyses between insulin levels and BW within each of the six studies with available insulin data (Table 3) and in a pooled analysis of all studies (Figure 3). While a strong correlation was observed in one study with fructose-fed rats (r^2^ = 0.5127), this was neither confirmed in another study in this model nor in both studies with ZSF1 rats or in two studies with HFD; of note, a numerically inverse correlation was observed in one study with HFD mice (see Online Supplement). In a pooled analysis of data from all animals in the hyperglycemic/diabetic groups, a week inverse correlation was observed (Figure 3, descriptive p-value 0.0094). Plasma insulin levels also positively correlated with BBW in the first fructose-feeding study but, if anything, inversely in the other five studies with available insulin data (Table 3).

**Table 3:** Correlation between plasma insulin and bladder and bladder/body weight acrossanimal models of T2DM. Animals from diabetic and non-diabetic group were pooled for each correlation analysis. Shown are total number of animals per model, squared correlation coefficient (r^2^) and descriptive p-value. *: negative slope (inverse correlation). A graphical representation of representative groups is shown in Figure 3, all other groups in the Online Supplement.

| n total | Bladder weight |  | Bladder/body weight |  |
| --- | --- | --- | --- | --- |
| | $r^2$ | p | $r^2$ | P |
| **ZSF1 rats (20-week, Hoechst)** |  |  |  |  |
| 30 | 0.0209 | 0.4461 | 0.0335* | 0.3329 |
| **ZSF1 rats (28-week, Hoechst)** |  |  |  |  |
| 45 | 0.0058* | 0.6192 | 0.0557* | 0.1186 |
| **Fructose-fed rats I (Mexico City)** |  |  |  |  |
| 12 | 0.5127 | 0.0088 | 0.4773 | 0.0129 |
| **Fructose-fed rats III (Mexico City)** |  |  |  |  |
| 14 | 0.0080 | 0.7605 | 0.1626* | 0.1529 |
| **HFD mice + semaglutide (Hoechst)** |  |  |  |  |
| 23 | 0.0529* | 0.2912 | 0.1046* | 0.1322 |
| **HFD mice (Mainz)** |  |  |  |  |
| 18 | 0.1319 | 0.2459 | 0.3389* | 0.0470 |

**Figure 3:**
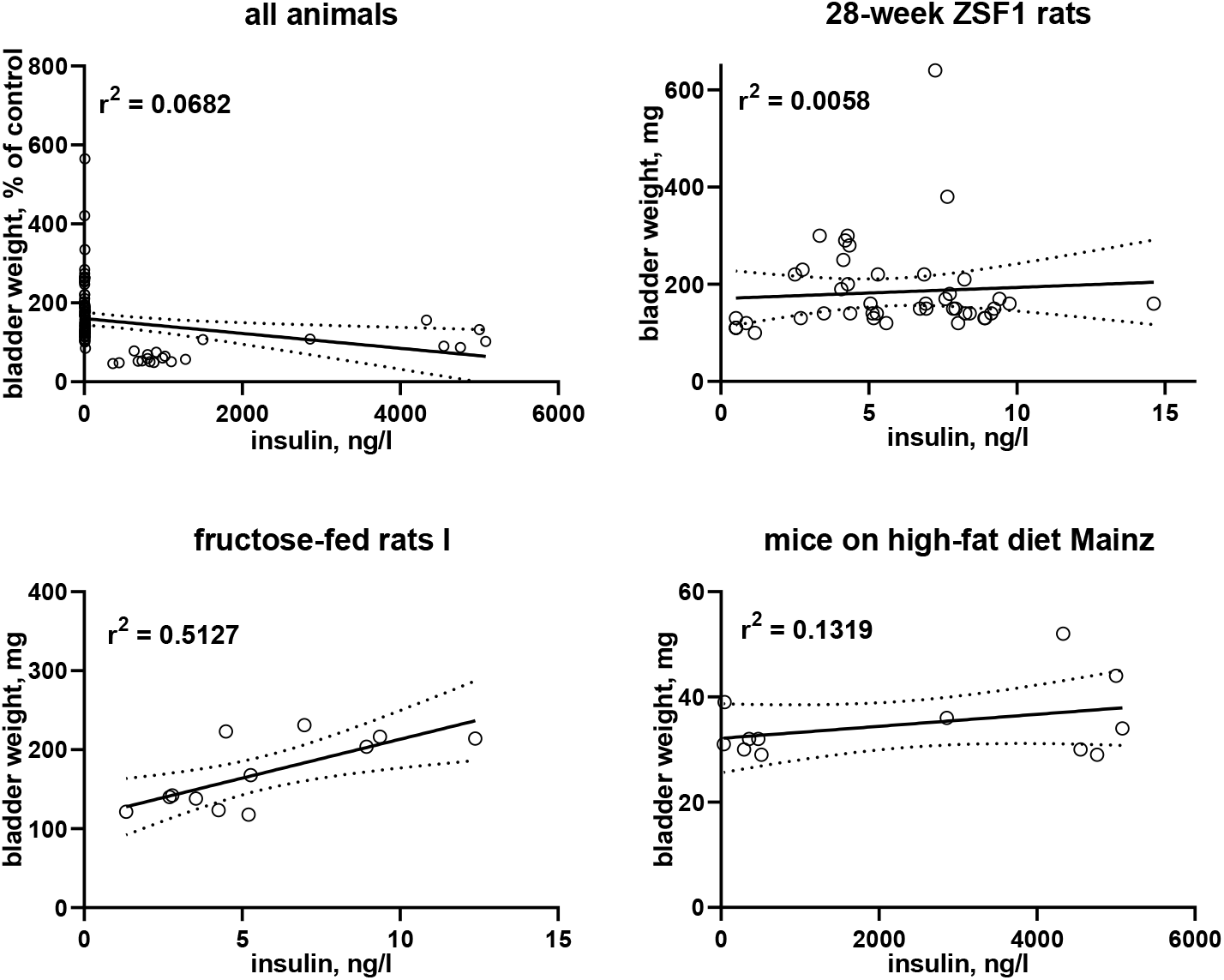
Correlation of bladder weight with insulin levels. To enable pooling of data from all studies, those for the upper left panel shows bladder weight only from the non-control groups expressed as % of mean values in the control group within a study. The other three panels show correlations within three representative studies; data from the remaining studies are shown in the Online Supplement. A quantitative description of the correlations is shown in Table 3. Mean values of bladder weight and insulin level in each study are shown in Table 1.

## Discussion

We have used data from 16 studies representing nine distinct rodent models of diabetes and 513 animals to address three specific questions:

- How widespread is urinary bladder enlargement in rodent models of experimental diabetes, particularly T2DM?
- How do diets and treatments other than insulin affect bladder enlargement?
- Is diabetic polyuria the key driver of diabetes-associated bladder enlargement?

### Critique of methods

It is a unique feature of the present study that it is fully based on data from experiments designed and conducted for other purposes. This is a limitation and also a strength. The limitation results from the fact that the original studies were neither designed nor powered to explore bladder enlargement and its causes; moreover, the 16 studies were heterogeneous in species (rat and mouse), T1DM vs. T2DM, specific aspects of models including genetic vs. acquired disease, duration of observation, and possible center differences between contributing laboratories. To accommodate this limitation, we have expressed data in the hyperglycemic/diabetic groups as % of the mean value in the corresponding euglycemic group for all inter-study analyses.

These limitations are outweighed by the use of an unprecedented number of models and studies. Given that each animal model of diabetes has limitations (19, 20), use of such variety of models should help to obtain data applicable to the heterogeneous population of diabetic patients (21). Moreover, using data from studies designed for other purposes fulfills the ethical mandate of the 3R principles to reduce the use of experimental animals wherever possible (22). Finally, it is highly cost-saving; generating a comparable number of models and studies for the primary purpose of the present analyses would have been too resource-intensive to be justifiable. Thus, the present analyses probably represent the largest collection of models and studies ever analyzed for any outcome parameter within a single project in diabetes research.

While it is customary to express organ enlargement relative to body weight, the body weight alterations in many diabetes models including those studied here (Table 1) would introduce a major bias in the analysis of bladder enlargement. Therefore, we have primarily focused on BW, but present BBW data as well for maximum transparency.

### Bladder enlargement across models

More than 70 previous studies have demonstrated a consistent enlargement of the urinary bladder in rats injected with STZ (mean BW 178% of control; range 99-440%) (9). A similar degree of enlargement was observed in a small number of studies with STZ-injected mice and rabbits, while other T1DM models including alloxan-injected rats and rabbits, BB/Wor rats and Akita mice exhibited a less pronounced increase in BW (10). Our studies with STZ-injected rats (two reported here, a third reported elsewhere (23)) confirm these findings. Moreover, we extend this to another model of T1DM, RIP-LCMV mice, for which no bladder weight data have been reported in the past.

We found in a previous systematic review that data on animal models of T2DM/obesity were much fewer in number and limited to five models: fructose-fed rats, HFD mice, Goto-Kakizaki rats, Zucker diabetic fatty rats and db/db mice (10). Across those models, bladder enlargement was small (about 150% of control) in fructose-fed rats and db/db mice, largely absent in HFD mice and in Goto-Kakizaki rats, but greater than the average enlargement in STZ-injected rats in Zucker diabetic fatty rats. Our present studies largely are in line with these findings. While we found a considerable enlargement in one study with fructose-fed rats, this was not confirmed in two other studies – by average confirming previous data in this model. In line with previous studies (10), we observed very minor if any bladder enlargement in db/db and in HFD mice. Our experiments also add data on four T2DM/obesity models for which bladder data had not been reported previously. We found a major increase in ZSF1 rats (>200% of control); as ZSF1 rats are a cross between Zucker diabetic fatty and spontaneously hypertensive rats and as Zucker rats were reported to exhibit a major bladder enlargement (9), these data are in line with previous findings. A moderate increase in bladder size was observed in rats injected with STZ at the neonatal stage and in ob/ob mice, whereas IRS2 knock-out mice did not exhibit bladder enlargement. In conclusion, the present data almost double the number of models of T2DM for which bladder size data have been reported. While previously reported (9, 10) and present data indicate that all animal models of T1DM exhibit bladder enlargement, although perhaps to a different extent, BW increases markedly in some models of T2DM, only moderately in others and not at all in additional models. Apparently, severity of diabetes as assessed by blood glucose levels does not explain the observed heterogeneity of bladder enlargement (see below). While the reasons for this heterogeneity are not fully clear, it is interesting that subgroups of patients with T2DM exhibiting distinct phenotypes are now also being recognized (21).

Other than in diabetes, bladder enlargement occurs in many conditions in animal models and patients, including bladder outlet obstruction and bladder denervation (24). It typically is associated with lower urinary tract dysfunction. Therefore, a better understanding of the pathophysiology underlying diabetes-associated bladder enlargement may help to define innovative treatment strategies to combat frequent lower urinary tract dysfunction in diabetic patients.

### Differential effects of diets and pharmacological treatments

The underlying studies had used various diets that had differential effects on glucose levels. While HFD increased glucose levels in most studies, canoletta and AMLN lowered it, whereas 0% choline/0.2% methionine did not affect it. As in previous studies (10), HFD was not associated with bladder enlargement in 3 present studies in mice. However, HFD was associated with bladder enlargement in a rat study (25). Diets were associated with congruent effects on glucose and BW in some cases such as canoletta, but observed effects differed in others: a diet containing 0% choline/0.2% methionine increased BW despite not affecting glucose, and AMLN did not affect BW despite lowering glucose.

The present studies are the first to explore effects of drug treatments other than insulin (10) on diabetes-associated bladder enlargement. The four drugs applied in the underlying studies had the expected effects or lack thereof on glucose levels for the model in which they were used. Like the diets, the pharmacological treatments did not affect glucose and BW in the same way in several cases: empagliflozin (a glucosuric drug (26)) lowered glucose but, if anything, increased BW; linagliptin (a drug not affecting glucosuria) tested within the same study caused a similar extent of bladder enlargement without affecting glucose levels. Semaglutide lowered glucose without affecting BW, and elafibranor did not affect glucose but reduced BW. These differential effects of diets and drug treatments are not easy to interpret because none of the studies had been designed to compare diet or drug effects on glucose and BW and because CI were wide in several cases. Nonetheless, the divergent effects casted doubt on the assumption that diabetic polyuria is the main reason for bladder enlargement.

### Role of glucose and insulin in bladder enlargement

When blood glucose levels exceed the renal reabsorption threshold of about 9-10 mM, the excreted glucose can act as an osmotic diuretic and cause diabetes-associated polyuria. It had been proposed that such polyuria is the main cause for bladder enlargement in experimental diabetes. Support for this hypothesis largely comes from the consistent observation that feeding with sucrose causes a similar degree of diuresis as STZ injection and a similar degree of bladder enlargement, while not affecting glucose levels to a major extent (11–17). Of note, these studies had used only one animal model of diabetes, STZ-injected rats. The polyuria hypothesis mechanistically implies that the degree of enlargement should be correlated with blood glucose levels because glucose levels determine the extent of diabetic polyuria, at least in models where glucose levels exceed the renal reabsorption threshold. The presence of bladder enlargement segregated only poorly with glucose levels relative to the renal reabsorption threshold in our analyses of 16 studies: small or no bladder enlargement was observed in two of 8 studies with glucose level >10 mM, whereas it was observed in one out of three studies with a glucose level < 9 mM; among the five studies with ambiguous glucose levels (9-10 mM), two exhibited bladder enlargement while three did not.

To further test the diabetic polyuria hypothesis, we have previously correlated the reported glucose levels and bladder size alterations at the group level across a total of >100 studies: while we detected a correlation at the group level, it was only of moderate strength, i.e., less than 20% in variability of BW could mathematically be attributed to that of glucose levels based on an r^2^ of 0.1772 (10). A major limitation of that analysis was that we only had access to data at the group level. We performed a similar correlation analysis based on individual animal data for glucose level and BW in a recent pilot study, which also yielded a correlation of only moderate strength (r^2^ 0.2013) (23). Therefore, individual animal-based correlation analyses were also performed for the 16 studies reported here as a pre-specified outcome parameter (Table 2). BW was correlated with blood glucose concentration in the three studies with T1DM models but only in three out of 13 studies in T2DM/obesity models. Moreover, the strength of correlation varied markedly across models. Thus, a strong correlation was observed in RIP-LCMV mice (r^2^ 0.723), a moderate correlation in STZ-injected rats, ZSF1 rats and rats with neonatal STZ injection (r^2^ 0.2346 - 0.4127), but correlations were very weak if existing at all in the other models. To corroborate these findings, we also performed a correlation analysis based on pooled individual animals from the hyperglycemic/diabetic groups of all 16 studies, which yielded an r^2^ of 0.0621 (Figure 2). Taken together, these data do not support the hypothesis that polyuria is the main factor to explain diabetes-associated bladder enlargement.

Insulin is not only a hormone but also a growth factor (27), and fructose-fed rats often exhibit a greater increase in insulin than in glucose levels, possibly reflecting peripheral insulin resistance (28). After having noticed a moderate to strong correlation of bladder enlargement with insulin levels in one study with fructose-fed rats (r^2^ 0.5127), we performed a similar post-hoc analysis on the other five models with available insulin data: all five studies including another study in fructose-fed rats exhibited very weak correlations, and in one of them (Table 3) and in the pooled analysis of all studies numerically inverse correlations were observed. This is not too surprising given that T1DM is characterized by a reduced presence of insulin; while insulin can be increased in models of T2DM including those reported here, this effect typically is counterbalanced by a reduced insulin sensitivity.

Thus, our data on diets, drug treatments, and most importantly our correlations between glucose and BW at the individual animal level do not support the diabetic polyuria hypothesis of bladder enlargement in animal models of T2DM. While this mechanism may play a role in some models such as RIP-LCMV mice, and perhaps a more moderate one in STZ-injected rats, it plays only a very minor if any role in most other models. While a role of insulin as mediator of bladder enlargement in diabetes would have been counter-intuitive based on reduced levels in T1DM and insulin resistance in T2DM, the lack of correlation between insulin levels and bladder size in all but one study provides additional experimental evidence in this regard. More generally, our data suggest that animal models of diabetes do not only differ in the presence and extent of bladder enlargement, but also in the pathophysiology leading to such enlargement in the models where it occurs. This conclusion is in line with the proposal that human T2DM is a heterogeneous condition with multiple underlying subgroups (21).

### Conclusions

Based on an unprecedented number of studies and animal models, we have shown that bladder enlargement is ubiquitous in animal models of T1DM and common, but not consistently present in those of T2DM/obesity. This heterogeneity among T2DM models is not explained by the severity of diabetes/hyperglycemia in those models, specifically not by glucose levels relative to the renal reabsorption threshold. For the first time we have explored effects of various diets and drug treatments other than insulin on diabetes-associated bladder enlargement; many of them had differential effects on glucose levels and bladder enlargement. These differential effects together with the generally moderate to absent association of glucose levels with BW do not support the hypothesis that diabetic polyuria is the main cause of diabetes-associated bladder enlargements – at least in most models. Further dedicated studies on bladder function relative to enlargement are needed. In a broader vein, our analyses highlight the heterogeneity between animal models of diabetes. While T2DM patients apparently also are a heterogeneous group (21), specific links between such subgroups and specific animal models remain to be established. Finally, our data demonstrate that major research accomplishments can be made without use of extra animals if smart planning is applied.

## Material and Methods

### Animal models

To collect information from a wide range of rodent models of diabetes in the spirit of the 3R principles (22), the present study is based on data from ongoing studies designed for other purposes; primary outcomes of these studies will be reported elsewhere by the respective investigators. In total, data were analyzed from 16 studies with 2-8 arms each representing 9 distinct T1DM and T2DM rat and mouse models (513 animals total). Details of each model are provided in the Online Supplement. Each of the underlying studies had been approved by the applicable independent committee or government agency for use and protection of experimental animals, and all studies were in line with the NIH guidelines for care and use of experimental animals (for details see Online Supplement). In each study, blood glucose concentration, body weight and BW were determined at study end in each animal and BBW was calculated. In some studies plasma insulin levels were also available. The following models were analyzed:

T1DM models: Two studies were done in male rats injected with STZ; one of them with additional treatment with the SGLT2 inhibitor empagliflozin (26) or the dipeptidylpeptidase 4 inhibitor linagliptin (29) in both control and STZ rats; the STZ-injected rats were considered as positive control because more than 70 studies have previously shown bladder enlargement in this model (9). As additional T1DM model we used male and female RIP-LCMV mice (30).

T2DM models: 13 studies representing 7 distinct models of T2DM were analyzed. Two studies were performed in male ZSF1 rats, a cross between a female Zucker Diabetic Fatty and a male spontaneously hypertensive SHHF rat (31), as compared to lean ZSF1 rats being observed up to an age of 20 or 28 weeks; each study included multiple treatment arms such as canoletta (hydrogenated rapeseed oil), 0% choline/0.2% methionine, AMLN diet (32), or the PPAR-α/δ agonist elafibranor (33). Three studies were performed in male fructose-fed rats (34) observed for 16 or 20 weeks. One study was done in male rats with STZ injection at the neonatal stage (35). One explored male and female IRS2 knock-out (36, 37) as compared to wild-type mice. Two studies looked at male and female ob/ob mice (38), one of them also including an arm with db/db mice of both sexes (38). Finally, three studies administered an HFD in male and female mice, one of them having an arm with treatment with the glucagon-like peptide-1 analog semaglutide (39).

### Data quality measures, data analysis and statistics

Except for studies performed in Ankara and Hoechst, the underlying studies did not involve randomization and/or blinding based on their specific needs; for details see Online Supplement. The decision to collect data on bladder weight from a given study was made before any animal was sacrificed. We operationally defined normoglycemia as blood glucose concentrations of <8 mM, hyperglycemia as 8-16 mM, and overt diabetes as >16 mM.

Sample size for the present analyses had been specified before data were viewed as all available animals from a given study. All analyses had been specified prior to data collection with one exception: after one study had included insulin data (fructose-fed I), additional correlation analysis was done for that parameter; when this yielded a correlation strength of likely biological relevance, presence of insulin data was checked for each study and, where available, correlations of bladder phenotype with insulin levels were performed.

The following pre-specified analyses were done for each study: The primary outcome parameter within each study was BW, analyzed as difference between the main hyperglycemic/diabetic and its control group with its 95% CI as derived from an unpaired, two-tailed t-test assuming comparable variability in both groups. The key secondary outcome parameter within each study was the correlation between blood glucose level and BW based on individual animal data of all groups with strengths of correlation assessed as the square of the correlation coefficient (r^2^) with its associated descriptive p-value. Other secondary outcome parameters were within-study group differences and correlations based on BBW and similar correlations with plasma insulin levels.

To explore correlations across groups, BW and BBW data from all animals other than those in the primary control group were expressed as % of the mean of the corresponding control group. This was followed by correlation analysis based on individual animal data across all models for comparison of BW vs. glucose, BBW vs. glucose, BW vs. insulin and BBW vs. insulin.

In line with recent guidelines and recommendations (40, 41), we consider all analyses reported here as exploratory. Therefore, no hypothesis-testing statistical analysis was applied and reported p-values should be considered descriptive and not hypothesis-testing. All calculations were performed using Prism (v9.02; GraphPad, Los Angeles, CA, USA).

### Study approval

All studies were in line with applicable rules and regulations including Directive 2010/63/EU of the European Parliament on the protection of animals used for scientific purposes and NIH guidelines for the use of experimental animals. The names of the responsible ethical animal committees and approval numbers for each of the 16 studies are provided in the Online Supplement.

## Author contributions

ZEY: experimentation for STZ study Ankara; overall data analysis; co-development of primary manuscript draft; reading and approval of final manuscript

JM: collection of data and supervision of study with ob/ob and IRS2 knock-out mice; reading and approval of final manuscript

EH: experimentation for RIP-LCMV mouse study; reading and approval of final manuscript

TRC: planning and experimentation for studies in ZSF1 rats and C57BL/6J, C57BL/6N, db/db, ob/ob and HFD mice; reading and approval of final manuscript

RE: planning and experimentation in C57BL/6N and HFD mice; reading and approval of final manuscript

JHBO: experimentation for fructose-fed and STZ rat studies; reading and approval of final manuscript

DLSV: experimentation for neonatal STZ rat study; reading and approval of final manuscript

NX: data collection and study supervision of the HFD model (Mainz); reading and approval of final manuscript

AK: planning of studies in in ZSF1 rats and C57BL/6J, C57BL/6N, db/db, ob/ob and HFD mice, reading and approval of final manuscript

UC: experimentation for RIP-LCMV mouse study; reading and approval of final manuscript

DC: supervision of fructose-fed, STZ and neonatal STZ rat studies; reading and approval of final manuscript

HL: study supervision of the HFD model (Mainz); reviewing and editing the manuscript; reading and approval of final manuscript

AP: conceptualization of overall project; reading and approval of final manuscript

EAI: supervision of STZ study Ankara; co-lead of overall project; reading and approval of final manuscript

MCM: conceptualization and lead of overall project; supervision of data analysis; co-development of primary manuscript draft; reading and approval of final manuscript

## Acknowledgments

This work was funded in part by TÜBITAK 2211/A (to ZEY), TÜBITAK-SBAG 118S443 and 119S769 (to EAI), Landesoffensive zur Entwicklung wissenschaftlich-ökonomischer Exzellenz (LOEWE; LOEWE Center for Translational Medicine and Pharmacology) of the State of Hessen, Germany (to UC), Conacyt Mexico 252702 (to DC), and Deutsche Forschungsgemeinschaft XI 139/2-1 (to NX), LI-1042/5-1 (to HL) and Mi 294/10-1 (to MCM). Some underlying studies were performed or funded by Sanofi-Aventis (identified as “Hoechst”) for purposes unrelated to this manuscript.

## Abbreviations

AMLN: Amylin liver NASH diet
BBW: bladder/body weight
BW: bladder weight
CI: confidence interval
HFD: high-fat diet
IRS2: insulin receptor substrate 2
RIP-LCMV: rat insulin promotor lymphocytic choriomeningitis virus
SGLT2: sodium-glucose transporter 2
STZ: streptozotocin
T1DM: type 1 diabetes mellitus
T2DM: type 2 diabetes mellitus
ZSF1: Zucker diabetic fatty/spontaneously hypertensive

