## Supplemental Information for "Urinary bladder enlargement across nine rodent models of diabetes: correlations with glucose and insulin levels"

All animals and data have been obtained from ongoing studies primarily performed for other purposes in the labs of the investigators. All studies were in line with applicable rules and regulations including Directive 2010/63/EU of the European Parliament on the protection of animals used for scientific purposes and NIH guidelines for the use of experimental animals. Data related to the primary purposes of those studies will be reported elsewhere. This Online Supplement includes the following information for each of the 16 studies:

- A description of each animal model
- Information on approval by the applicable animal committee
- A graphical depiction of values for each animal within each study
- A graphical depiction of correlations between glucose and insulin levels on the one and bladder and bladder/body weight data within each study. Outcomes of correlation analyses are shown as $r^2$ values to indicate strength of correlation, irrespective whether the correlations had descriptive p-values < or ≥ 0.05.

#### Type 1 diabetes models

##### STZ-injected rats (Mexico City)

The study had been approved by Institutional Ethics Committee (Cicual-Cinvestav; 0102-14). This was a 2-armed study: Male Wistar rats (250 g, 7 weeks old) were obtained from the laboratory animal facility of the Dept. of Pharmacobiology, Vinvestag, Sede Sur and injected i.p. with vehicle (citrate buffer pH 4.5) or 60 mg/kg streptozotocin (STZ). After 6 weeks, they were anesthetized with isoflurane (3%) and euthanized by decapitation.

Supplemental Figure 1: Blood glucose, body weight, bladder weight and bladder/body weight in STZ-injected rats. Each data point represents one animal, bars and error bars represent means ± SD.

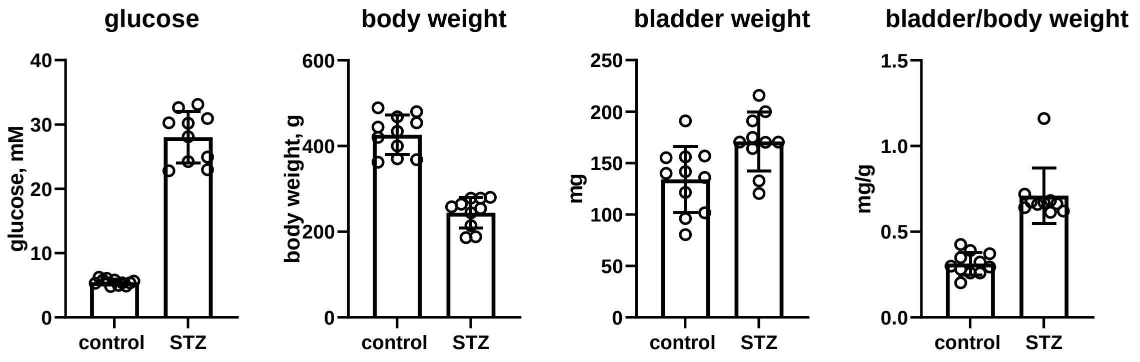

Supplemental Figure 2: Correlation of blood glucose with bladder and bladder/body weight. Each data point represents one animal; the line represents the calculated regression line with its 95% CI. Descriptive p-values were 0.0261 and <0.0001, respectively.

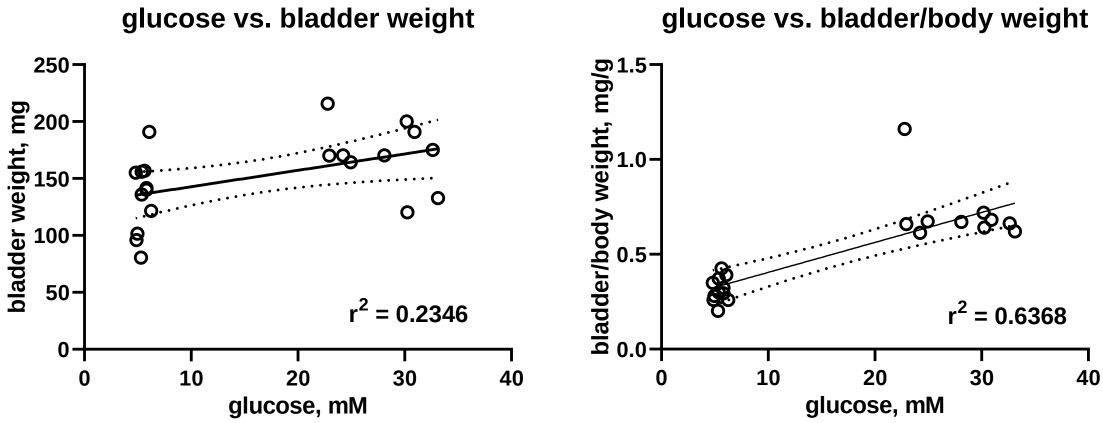


#### STZ-injected rats (Ankara)

The study had been approved by the animal welfare committee of Ankara University (2019-4-41). In a 6-armed study, 11-week-old male Sprague Dawley rats were obtained from Bilkent University Genetics and Biotechnology Research Center and Kobay Experimental Animals Laboratory (Ankara, Turkey) and injected with vehicle or 50 mg/kg, i.p. STZ; within each group, animals were further subdivided after 12-15 weeks by allocating them to treatment with vehicle, empagliflozin (30 mg/kg, with oral gavage) or linagliptin (4 mg/kg, with oral gavage). After 8-10 weeks of treatment, they were anesthetized under 2% isoflurane inhalation and sacrificed. Rats were randomized to group allocation and the person obtaining and weighing tissues was blinded to group allocation. Glucose levels had been measured in the week preceding sacrificing the rats.

Supplemental Figure 3: Blood glucose, body weight, bladder weight and bladder/body weight in STZ-injected rats with and without additional treatment with empagliflozin (empa) or linagliptin (lina). Each data point represents one animal, bars and error bars represent means ± SD. Note that measured glucose values were censored at 33.3 mM.

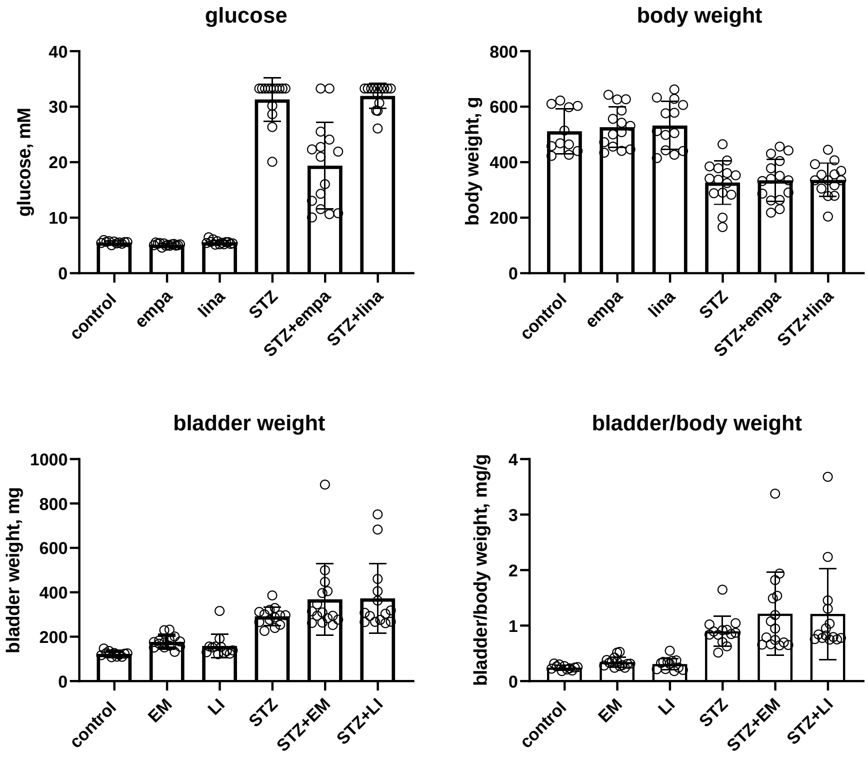

Supplemental Figure 4: Correlation of blood glucose with bladder and bladder/body weight. Each data point represents one animal; the line represents the calculated regression line with its 95% CI. Descriptive p-values were <0.0001 for both comparisons.

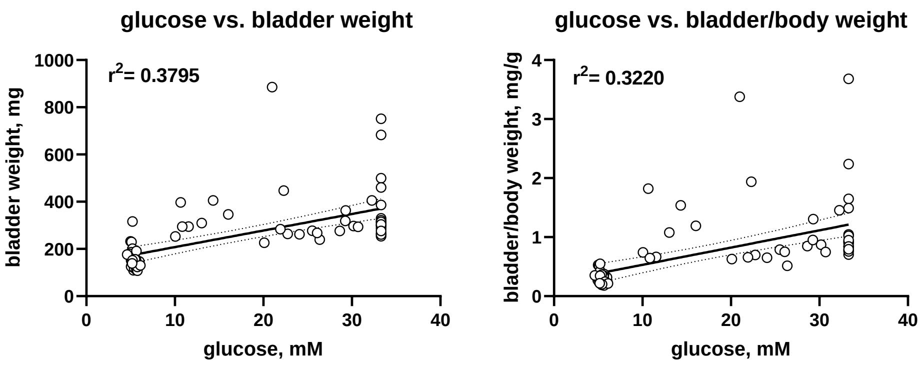


#### Rat insulin promotor lymphocytic choriomeningitis virus (RIP-LCMV) mice (Frankfurt)

RIP-LCMV-GP mice express the glycoprotein (GP) of the lymphocytic choriomeningitis virus (LCMV) under control of the rat insulin promoter (RIP) in the beta-cells of the islets of Langerhans (1). Infection of such mice with LCMV as environmental trigger initiates an immune response directed against LCMV as well as the transgenically expressed GP in the beta-cells resulting in type 1 diabetes (T1DM) within 10-14 days after infection (2). Generation and screening by PCR of RIP-LCMV-GP transgenic mice were as previously described (1). These mice have been backcrossed to a C57BL/6J background for more than 30 years. LCMV Armstrong clone 53b (LCMV-Arm) were plaque-purified three times on Vero cells and stocks were prepared by a single passage on BHK-21 cells (von Herrath et al., 1994 Immunity 1: 231-242).

The study had been approved by the Ethics Committee of the State Ministry of Agriculture, Nutrition and Forestry, State of Hessen, Germany (V54-19c20/15-FU-1192 and V54-19c20/15-FU-1213). This was a 2-armed study: 8–10-week-old female and male RIP-LCMV-GP mice (C57BL/6J background) were obtained from mfd-diagnostics (Wendelsheim, Germany). Initial generation and screening by PCR of RIP-LCMV-GP mice (H-$2^b$) were as previously described (3). To induce T1D, RIP-LCMV-GP mice were infected with 5 x $10^3$ plaque forming units (pfu; intraperitoneally in 100 µl RPMI medium). The majority (90-95%) of LCMV-infected RIP-LCMV-GP mice develop T1D within 10-14 days after infection (blood glucose levels >16 mM). Blood glucose measurements were performed using a SD code-free blood glucose monitoring system (SD Biosensor, South Korea). Uninfected, age-matched female and male RIP-LCMV-GP mice were used as control. Mice were sacrificed 6-8 weeks after LCMV-infection by isoflurane overdose and cervical dislocation. For the current experiments, bladders have been removed post-mortem from LCMV-infected, but otherwise untreated and from control mice involved in other projects. Bladders have been homogenized in 3 ml TriReagent (Sigma-Aldrich, St. Louis, MO), and stored at -80C until further use

Supplemental Figure 5: Blood glucose, body weight, bladder weight and bladder/body weight in RIP-LCMV mice. Each data point represents one animal, bars and error bars represent means ± SD. Note that measured glucose values were censored at 33.3 mM.

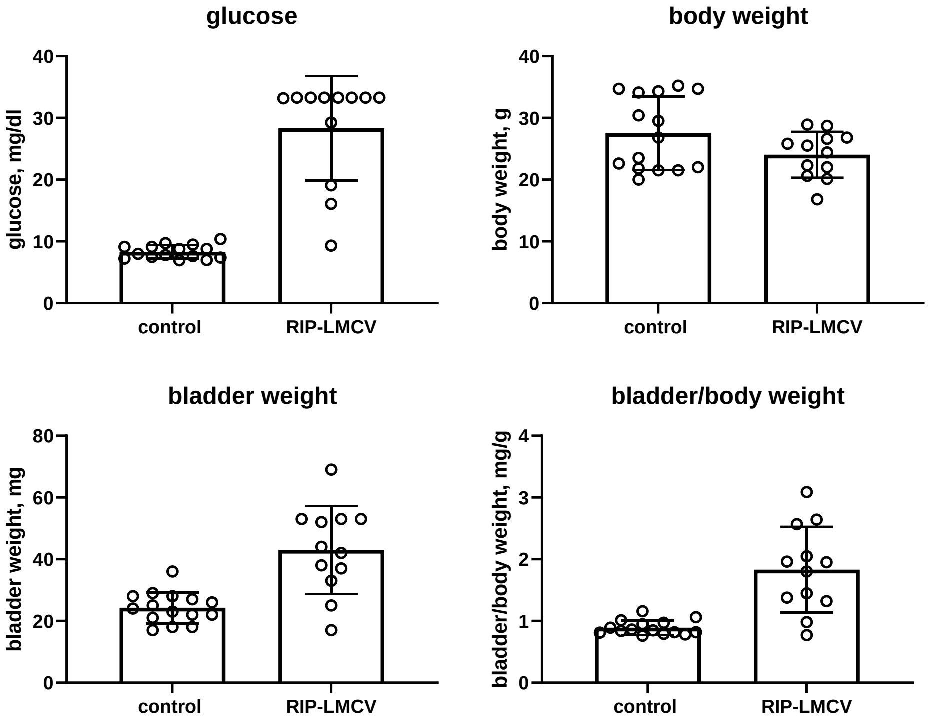

Supplemental Figure 6: Correlation of blood glucose with bladder and bladder/body weight in RIP-LMCV mice. Each data point represents one animal; the line represents the calculated regression line with its 95% CI. Descriptive p-values were <0.0001 in both cases.

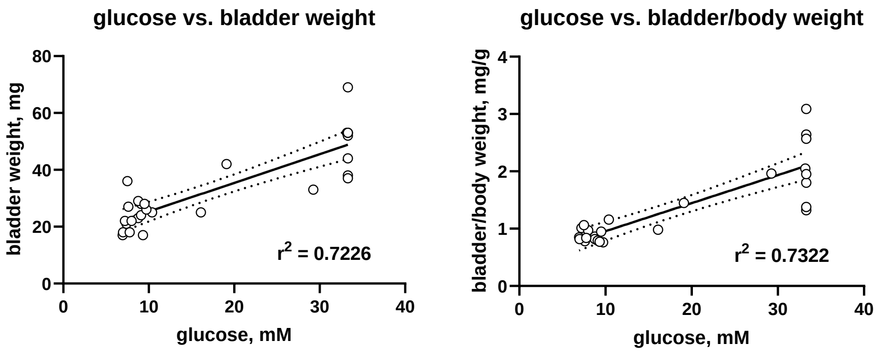


### Type 2 diabetes models

#### ZSF1 rats (Hoechst)

Two studies of similar design but with different treatment groups were performed in 7-8-week-old male ZSF1 rats (Charles River, Kingston, NY, USA). The first had a duration of 12 weeks (20 weeks old rats) and the second of 20 weeks (28 weeks old rats). Both studies had been approved by the Ethics Committee of the State Ministry of Agriculture, Nutrition and Forestry, State of Hessen, Germany (the V54 - 19 c 20/15-FH/Anz. 1012). Rats were randomized by body weight before allocation to the different diets (ssniff Spezialdiäten, GmbH, Soest, Germany). The 12-week study included 6 arms: lean ZSF1 control rats, obese ZSF1 rats, obese ZSF1 rats on AMLN diet (4), obese ZSF1 rats on an AMLN diet in which 15% primex had been replaced with canoletta, and obese ZSF1 rats on a 0% choline/0.2% methionine diet. The 20-week study had 8 arms: lean ZSF1 control rats, obese ZSF1 rats, obese ZSF1 rats on AMLN diet + 99.5% methylcellulos and 0.5% Tween-80 (vehicle for elafibranor), obese ZSF1 rats on AMLN diet + the PPAR-α/δ agonist elafibranor (30 mg/kg) (5), obese ZSF1 rats on 0% choline/0.2% methionine diet, obese ZSF1 rats on canoletta diet, obese ZSF1 rats on AMLN diet + oil (vehicle for $CCl_4$), and obese ZSF1 rats on AMLN diet + $CCl_4$ (0.2 mg/kg); of note, this dose of $CCl_4$ mistakenly was much lower than planned and not considered to cause hepatic cirrhosis. At the end of the study the rats were euthanized by final exsanguination under isoflurane anesthesia.

Supplemental Figure 7: Blood glucose, body weight, bladder weight and bladder/body weight in lean control and ZSF1 rats on standard diet or the indicated specific diets after 12 weeks. Each data point represents one animal, bars and error bars represent means ± SD.

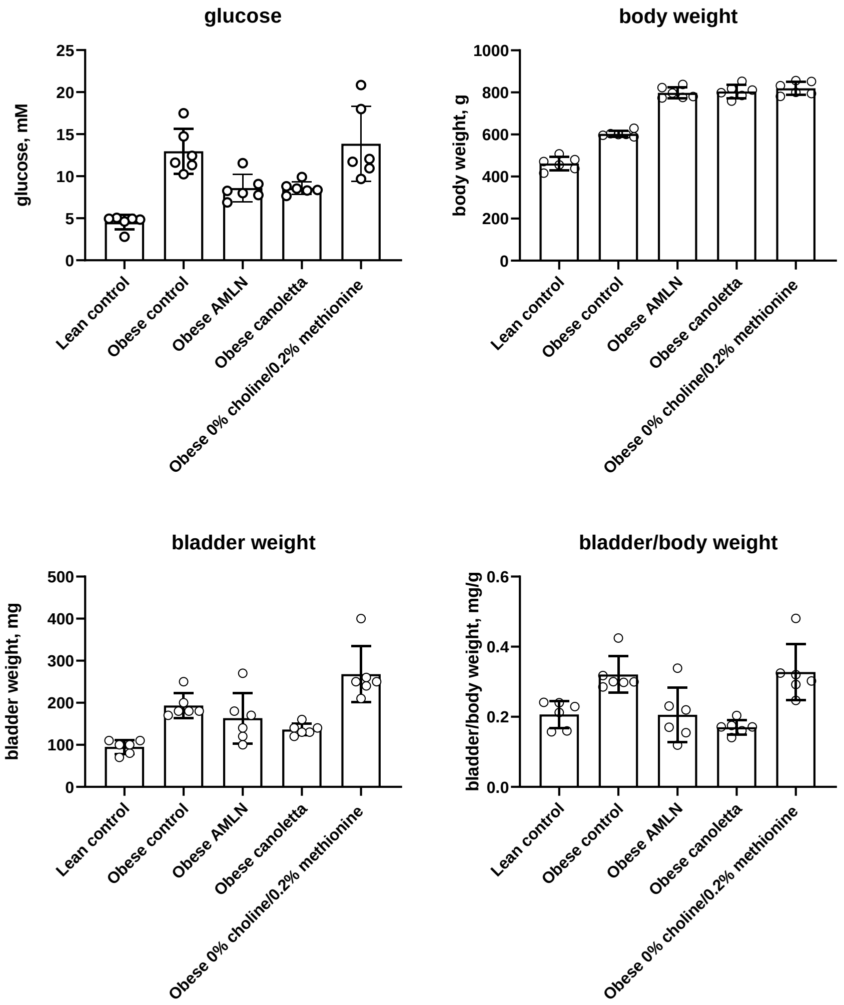

Supplemental Figure 8: Correlation of blood glucose (upper panels) and insulin (lower panels) with bladder and bladder/body weight in in lean control and ZSF1 rats with or without specific diets after 12 weeks. Each data point represents one animal; the line represents the calculated regression line with its 95% CI. Descriptive p-values were 0.4461, 0.3329, 0.6192 and 0.1186, respectively. Note that insulin levels in all lean control rats were below detection limit (0.512 ng/l) and were entered at this value into the correlation analyses.

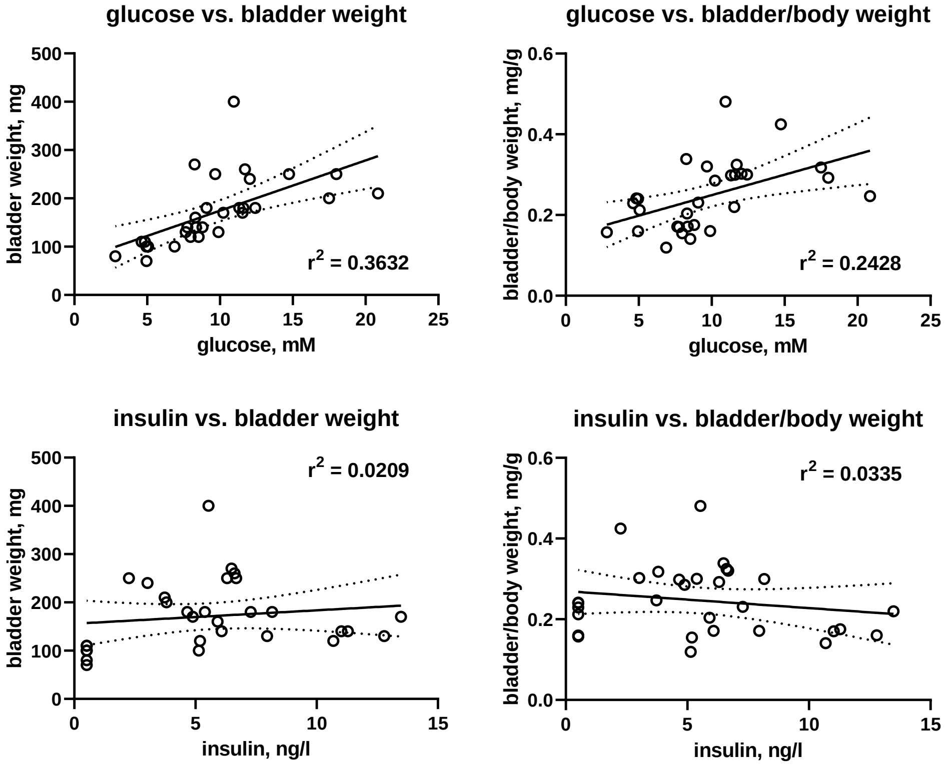

Supplemental Figure 9: Blood glucose, body weight, bladder weight and bladder/body weight in lean control and ZSF1 rats on standard diet or the indicated specific diets after 20 weeks. Each data point represents one animal, bars and error bars represent means ± SD.

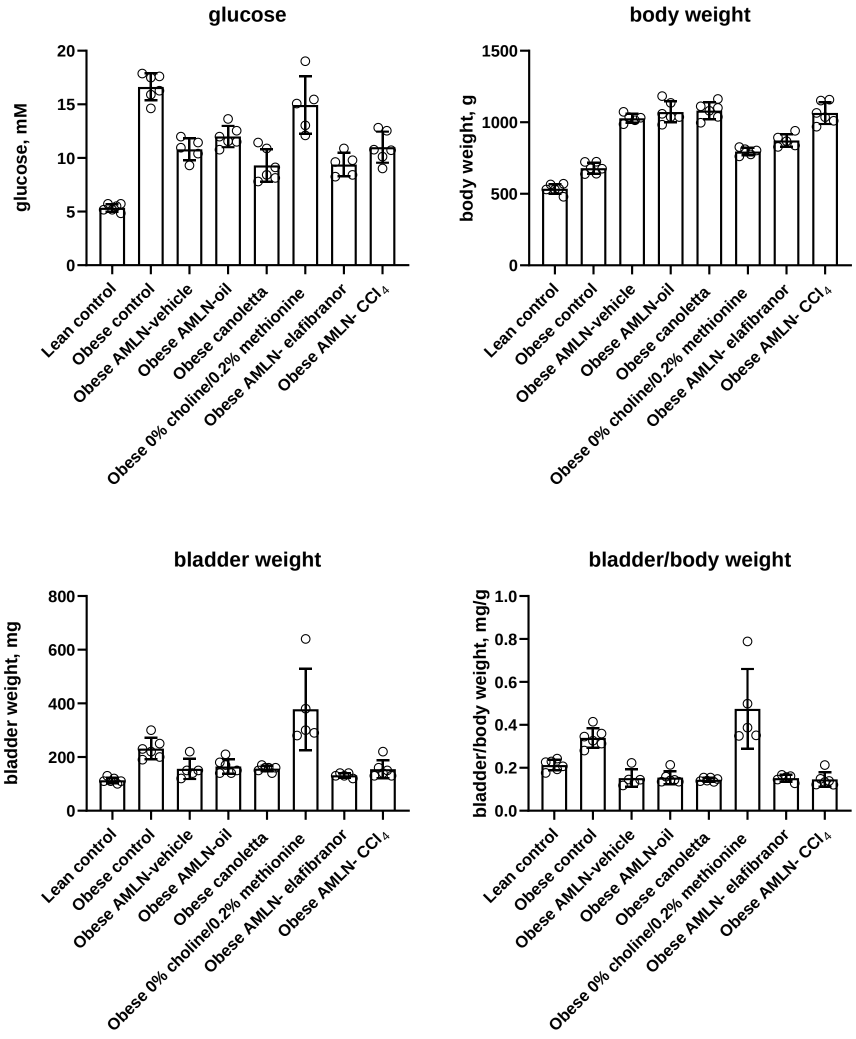

Supplemental Figure 10: Correlation of blood glucose (upper panels) and insulin (lower panels) with bladder and bladder/body weight in in lean control and ZSF1 rats with or without specific diets after 20 weeks. Each data point represents one animal; the line represents the calculated regression line with its 95% CI. Descriptive p-values were <0.0001, <0.0001, 0.6192 and 0.1186, respectively. Note that insulin levels in 4 lean control rats were below detection limit (0.512 ng/ml) and were entered at this value into the correlation analyses.

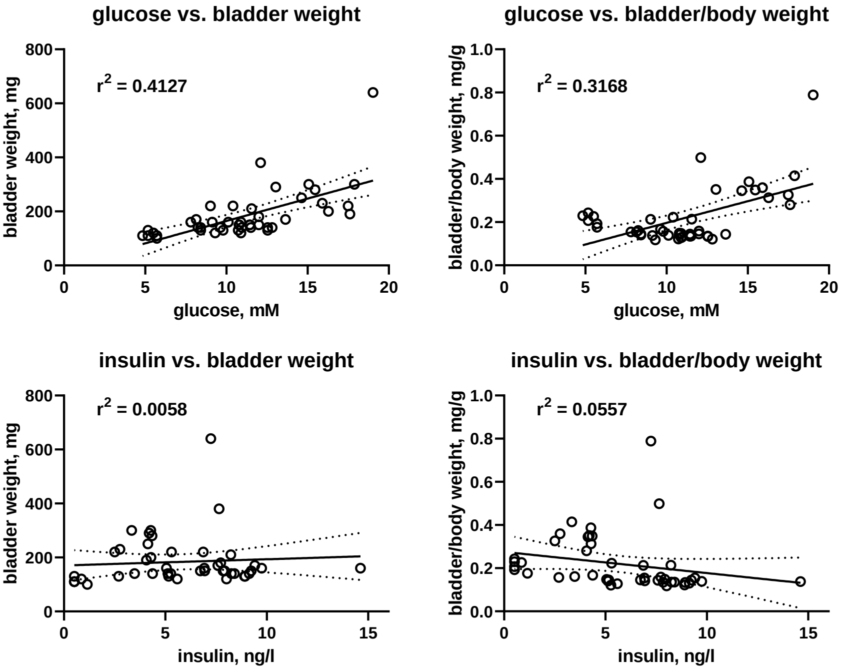


#### Fructose-fed rats (Mexico City)

Three studies of similar design but different observation periods were performed in fructose-fed rats (6). All three studies had been approved by the institutional ethics committee (Cicual-Cinvestav, approval 0102-14). They were 2-armed studies: Male Wistar rats (200-220 g, 7 weeks old) were obtained from the laboratory animal facility of the Dept. of Pharmacobiology, Cinvestav Sede Sur and fed a standard diet or a diet supplemented with a 15% fructose solution. After 16 weeks (study II) or 20 weeks (studies I and III), they were anesthetized with isoflurane (3%) and euthanized by decapitation.

Supplemental Figure 11: Blood glucose, body weight, bladder weight and bladder/body weight in study I of control and fructose-fed rats. Each data point represents one animal, bars and error bars represent means ± SD.

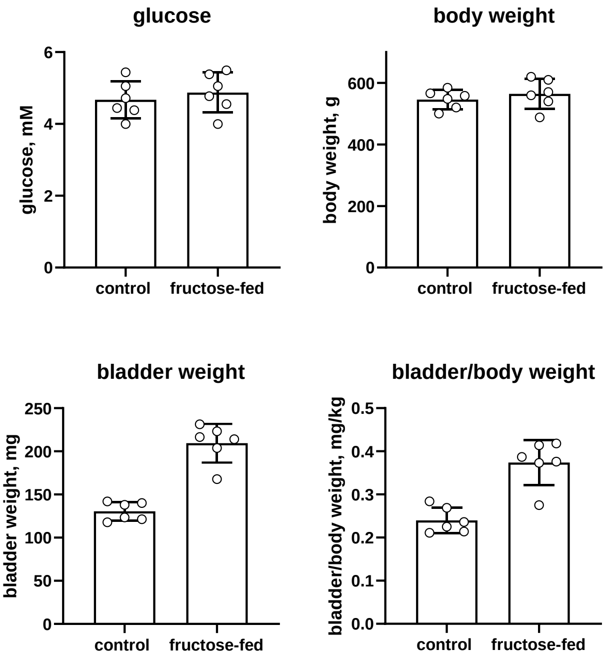

Supplemental Figure 12: Correlation of blood glucose (upper panels) or insulin (lower panels) with bladder and bladder/body weight in study I of control and fructose-fed rats. Each data point represents one animal; the line represents the calculated regression line with its 95% CI. Descriptive p-values were 0.7465, 0.8384, 0.0088 and 0.0129, respectively.

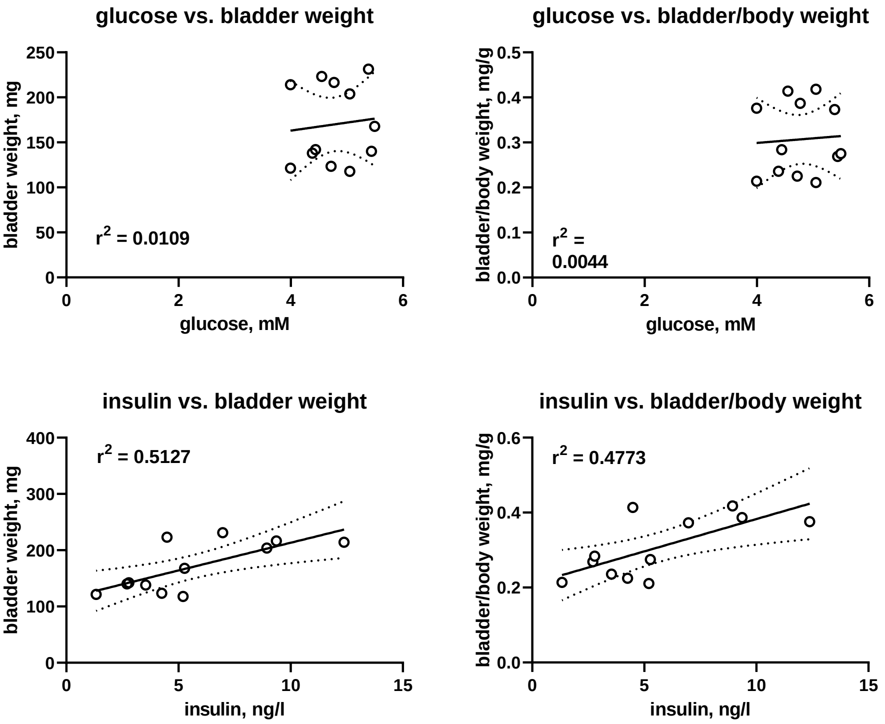

Supplemental Figure 13: Blood glucose, body weight, bladder weight and bladder/body weight in study II of control and fructose-fed rats. Each data point represents one animal, bars and error bars represent means ± SD.

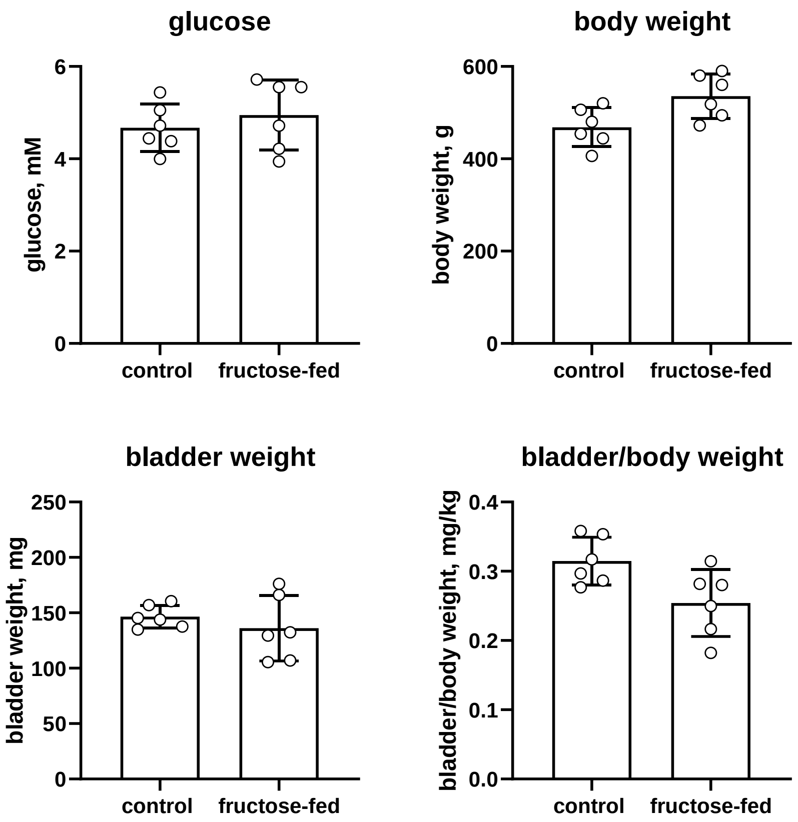

Supplemental Figure 14: Correlation of blood glucose with bladder and bladder/body weight in study II of control and fructose-fed rats. Each data point represents one animal; the line represents the calculated regression line with its 95% CI. Descriptive p-values were 0.1473 and 0.0944, respectively.

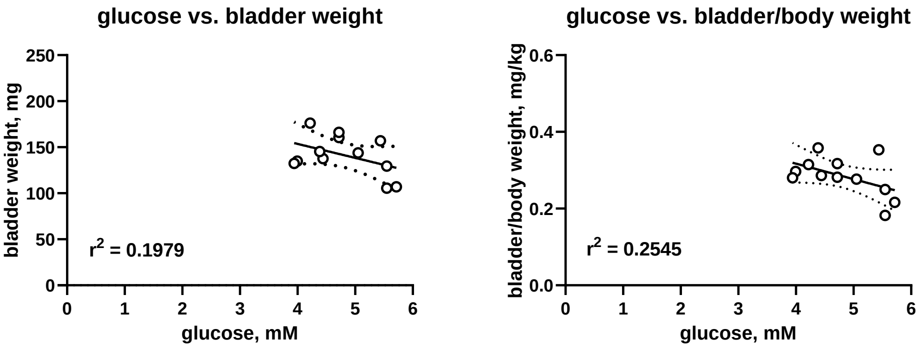

Supplemental Figure 15: Blood glucose, body weight, bladder weight and bladder/body weight in study III of control and fructose-fed rats. Each data point represents one animal, bars and error bars represent means ± SD.

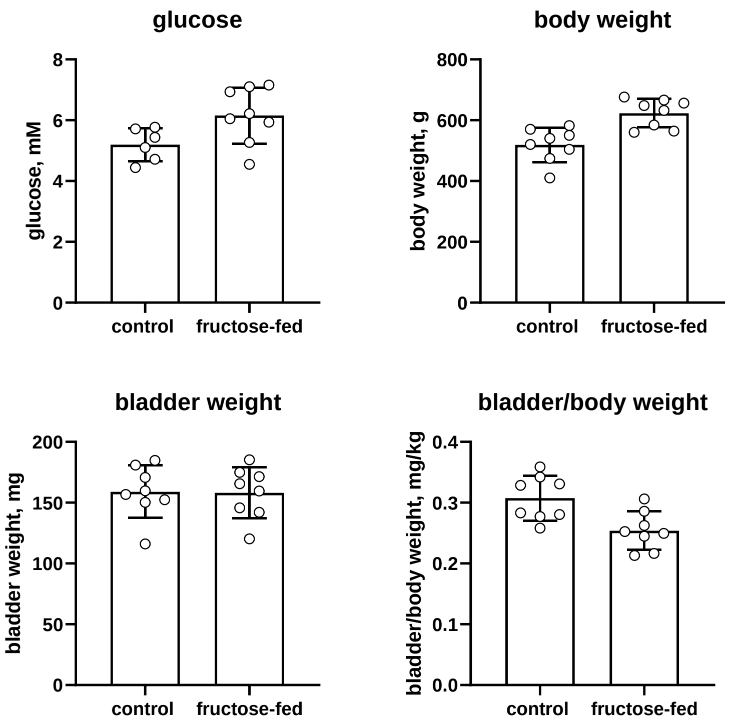

Supplemental Figure 16: Correlation of blood glucose and insulin with bladder and bladder/body weight in study III of control and fructose-fed rats. Each data point represents one animal; the line represents the calculated regression line with its 95% CI. Descriptive p-values were 0.4481, 0.4590, 0.7605 and 0.1529, respectively.

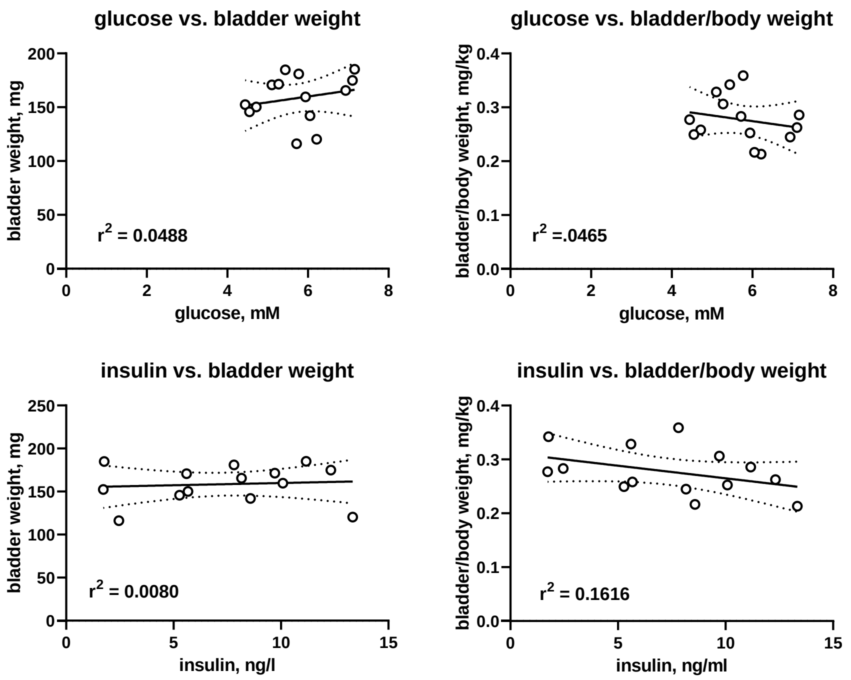


#### Rats with neonatal STZ injection (Mexico City)

The study had been approved by the institutional ethics committee (Cicual-Cinvestav, approval 0102-14). This was a 2-armed study: Male Wistar newborn rats (7-10 g, 3-5 days old) were obtained from the laboratory animal facility of the Dept. of Pharmacobiology, Cinvestav Sede Sur and injected i.p. with vehicle (citrate buffer pH 4.5) or 70 mg/kg STZ. After 16 weeks, they were anesthetized with isoflurane (3%) and euthanized by decapitation.

Supplemental Figure 17: Blood glucose, body weight, bladder weight and bladder/body weight in control animals and rats with neonatal STZ injection. Each data point represents one animal, bars and error bars represent means ± SD.

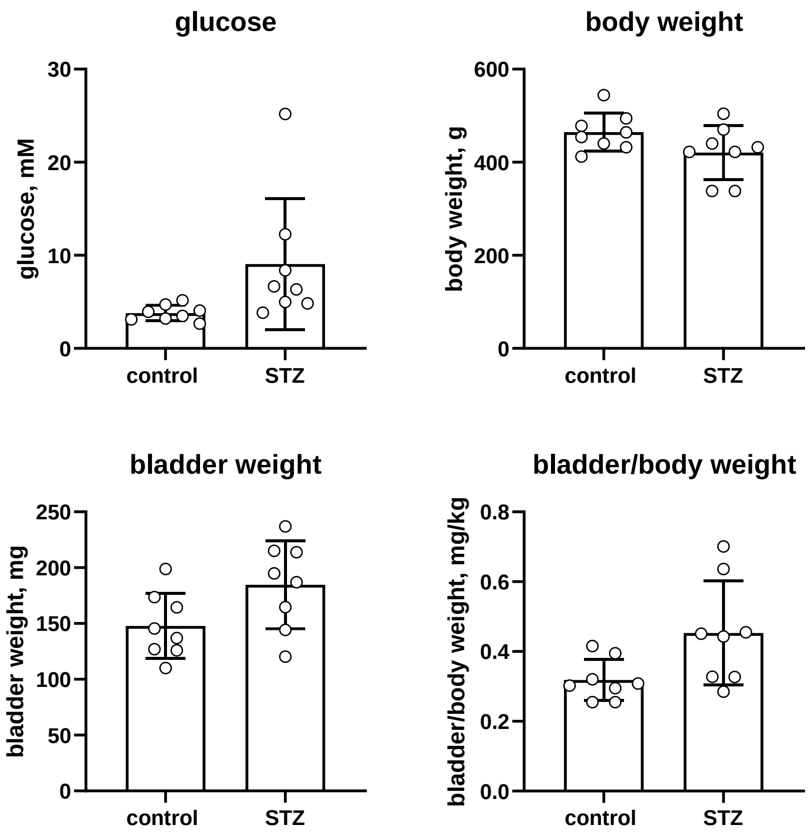

Supplemental Figure 18: Correlation of blood glucose with bladder and bladder/body weight in control animals and rats with neonatal STZ injection. Each data point represents one animal; the line represents the calculated regression line with its 95% CI. Descriptive p-values were 0.0199 and 0.0003, respectively.

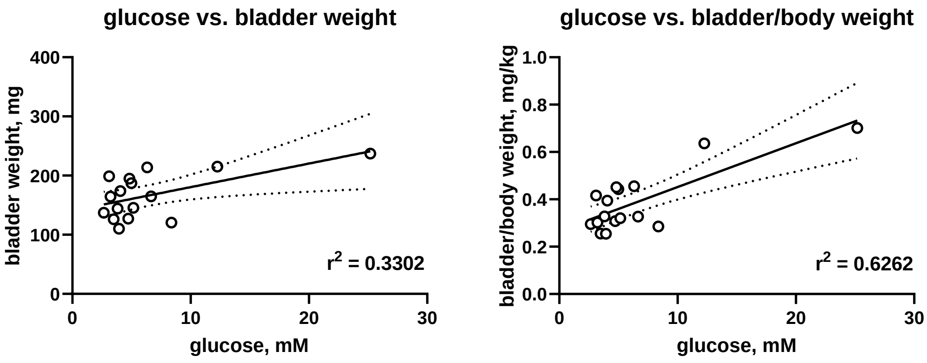


#### IRS2 knock-out mice (Cologne)

Animal breeding, maintenance and experiments had been approved by the responsible federal state authority (Landesamt für Natur-, Umwelt- und Verbraucherschutz Nordrhein-Westfalen; 84-02.04.2016.A049 and 84-02.04.2016.A422). This was a 2-armed study using mice from both sexes: Details of the IRS2 knock-out mouse model have been described (7, 8). Tail or ear clips from 3-week-old mice were processed for genotyping. Mice had an C57BL/6J background and were kept in individually ventilated cages with a 12h/12h dark/light cycle and food and water *ad libitum*. We used rat/mouse maintenance food (V1554-703, ssniff Spezialitäten GmbH, Soest, Germany). Immediately after killing by cervical dislocation, urinary bladder was excised and weighed and blood glucose was measured using a blood-glucose meter (Accu-Check® Aviva, Roche Diagnostics Deutschland GmbH, Mannheim, Germany) with a drop of blood leaking from the cut tail.

Supplemental Figure 19: Blood glucose, body weight, bladder weight and bladder/body weight in control (C57BL/6) and IRS2 knock-out mice. Each data point represents one animal, bars and error bars represent means ± SD.

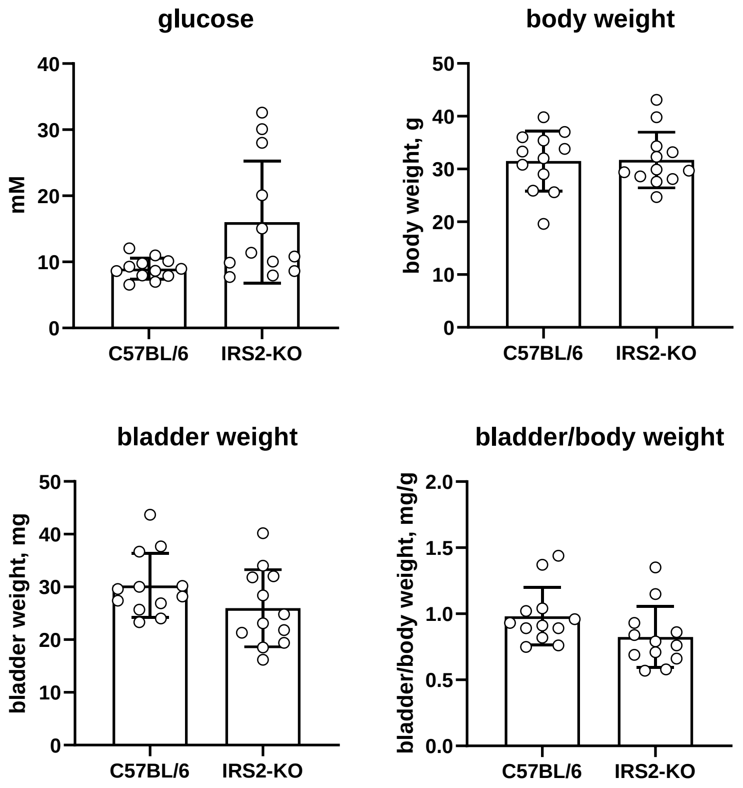

Supplemental Figure 20: Correlation of blood glucose with bladder and bladder/body weight in control (C57BL/6) and IRS2 knock-out mice. As the study had used mice aged 16-61 weeks, additional correlation analysis of age vs. bladder weight was performed to explore whether age differences may have affected outcomes; this has not been the case (lower panel). Each data point represents one animal; the line represents the calculated regression line with its 95% CI. Descriptive p-values were 0.0893, 0.1305 and 0.6776, respectively.

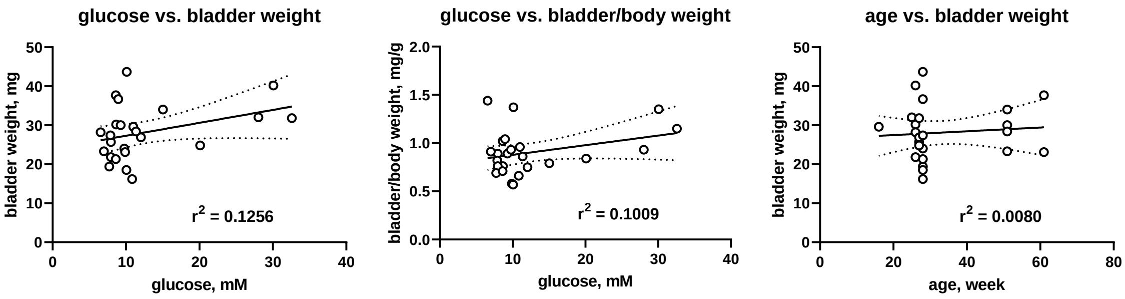


#### ob/ob mice (Cologne)

This study used control (C57BL/6J background) and ob/ob mice (leptin B6.Cg-Lep ob/J) from both sexes initially purchased from the Jackson Laboratory (Bar Harbor, USA) and maintained at our facility. Ethical approval, genotyping and study conduct were identical with those described above for the IRS2 knock-out experiments.

Supplemental Figure 21: Blood glucose, body weight, bladder weight and bladder/body weight in control (C57BL/6) and ob/ob mice. Each data point represents one animal, bars and error bars represent means ± SD.

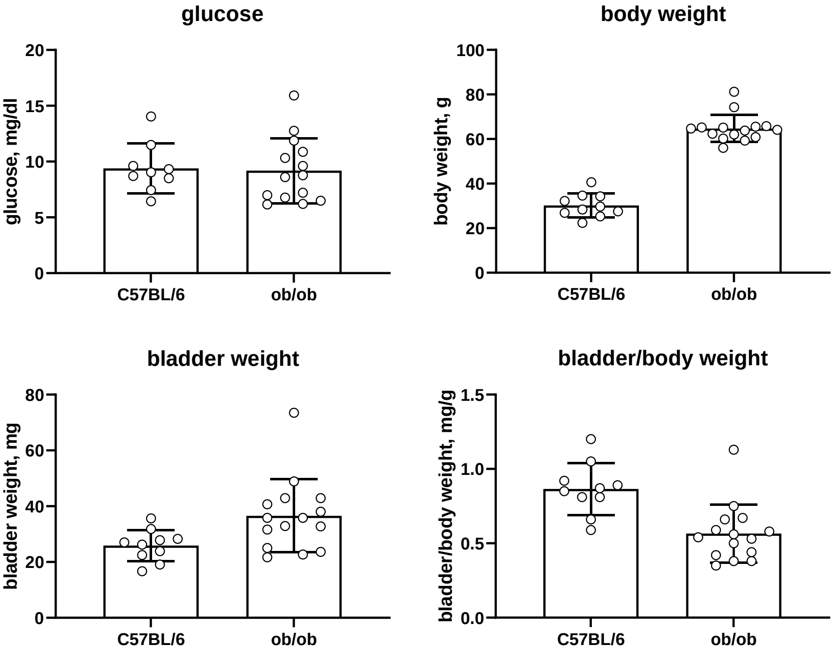

Supplemental Figure 22: Correlation of blood glucose with bladder and bladder/body weight in control (C57BL/6) and ob/ob mice. As the study had used mice aged 21-56 weeks, additional correlation analysis of age vs. bladder weight was performed to explore whether age differences may have affected outcomes; this has not been the case (lower panel). Each data point represents one animal; the line represents the calculated regression line with its 95% CI. Descriptive p-values were 0.7410, 0.9593 and 0.4126, respectively.

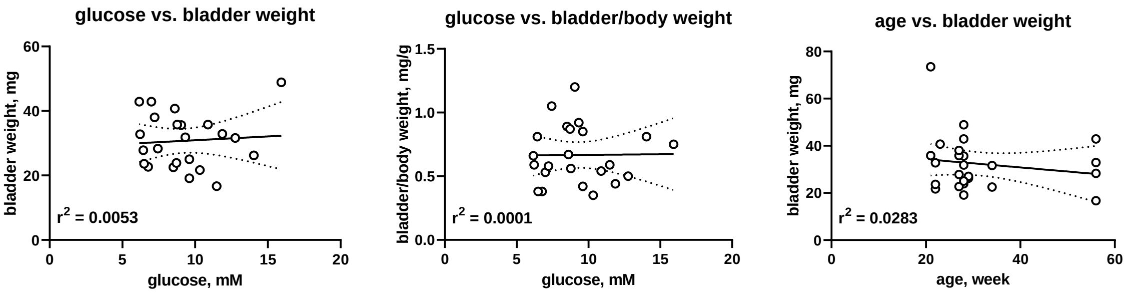


#### ob/ob and db/db mice (Hoechst)

This was a study in which ob/ob and db/db mice were studied in comparison to wild-type C57BL/6J mice. As the underlying study involved only tissue harvesting and no experimentation or other intervention in living animals, no animal permit from the authorities was required according to §4 Abs. 3 of the German animal protection law (Tierschutzgesetz). Instead an internal permit for animal numbers to be reported (Tiertötung einer Maus, T4-12.A) was obtained. Mice of both sexes were obtained from Charles River Laboratories Germany GmbH (Sulzfeld, Germany) for tissue collection at an age of 12 weeks. All mice were euthanized with an isoflurane overdose and euthanized by cervical dislocation, half within each group at 7 a.m. (fed stage) and half at 2 p.m. (starved stage). The graphs show pooled data of both sexes and both time points of euthanization.

Supplemental Figure 23: Blood glucose, body weight, bladder weight and bladder/body weight in control (C57BL/6J), db/db and ob/ob mice. All mice were euthanized with an isoflurane overdose and euthanized by cervical dislocation, half within each group at 7 a.m. (fed stage) and half at 2 p.m. (starved stage). The graphs show pooled data of both sexes and both time points of euthanization. Each data point represents one animal, bars and error bars represent means ± SD.

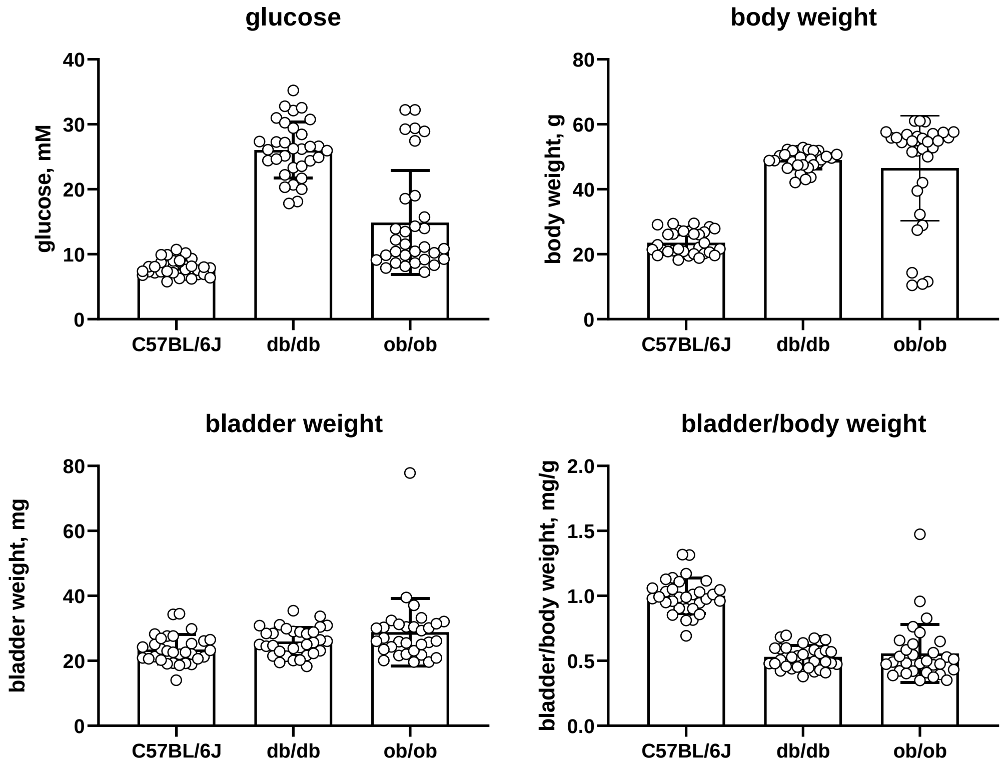

Supplemental Figure 24: Correlation of blood glucose with bladder and bladder/body weight in control (C57BL/6J), db/db and ob/ob mice. Each data point represents one animal; the line represents the calculated regression line with its 95% CI. Note that almost all animals in the upper left quarter of the glucose vs. bladder/body weight panel represent the control group driven by the lower body weight in absence of change in bladder weight in this group. Descriptive p-values were 0.2521 and <0.0001, respectively.

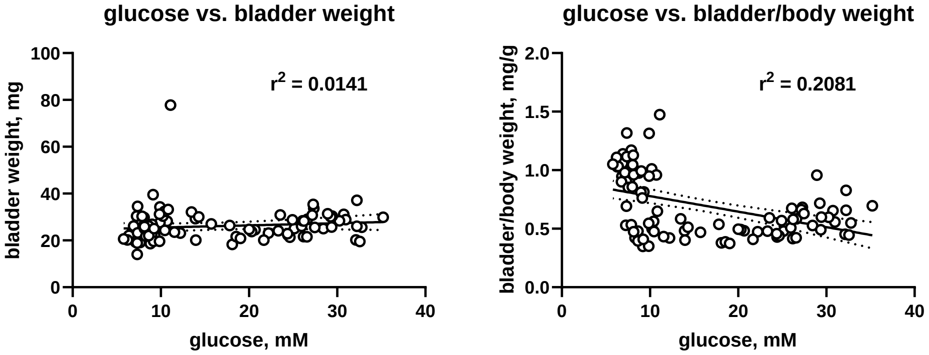


#### HFD mice (Hoechst)

This was a 2-armed study in 12-week-old C57BL/6N control and 24-week-old HFD mice, which was performed in conjunction with the above study on db/db and ob/ob mice and had been approved by the Ethics Committee of the State Ministry of Agriculture, Nutrition and Forestry, State of Hessen, Germany (the V54 - 19 c 20/15-FH/Anz. 1024 and T4-12.A1). Mice of both sexes were obtained from Charles River Laboratories Germanny GmbH (Sulfeld, Germany); the HFD group received an HFD (EF adjusted fat diet, 25% fat content; ssniff Spezialdiäten, GmbH, Soest, Germany) for 18 weeks for tissue collection at an age of 24 weeks. All mice were euthanized with an isoflurane overdose and euthanized by cervical dislocation, half within each group at 7 a.m. (fed stage) and half at 2 p.m. (“fasted” stage). The graphs show pooled data of both sexes and both time points of euthanization.

Supplemental Figure 25: Blood glucose, body weight, bladder weight and bladder/body weight in mice (C57BL/6N) on a control or an HFD. All mice were euthanized with an isoflurane overdose and euthanized by cervical dislocation, half within each group at 7 a.m. (fed stage) and half at 2 p.m. (“fasted” stage). Each data point represents one animal, bars and error bars represent means ± SD.

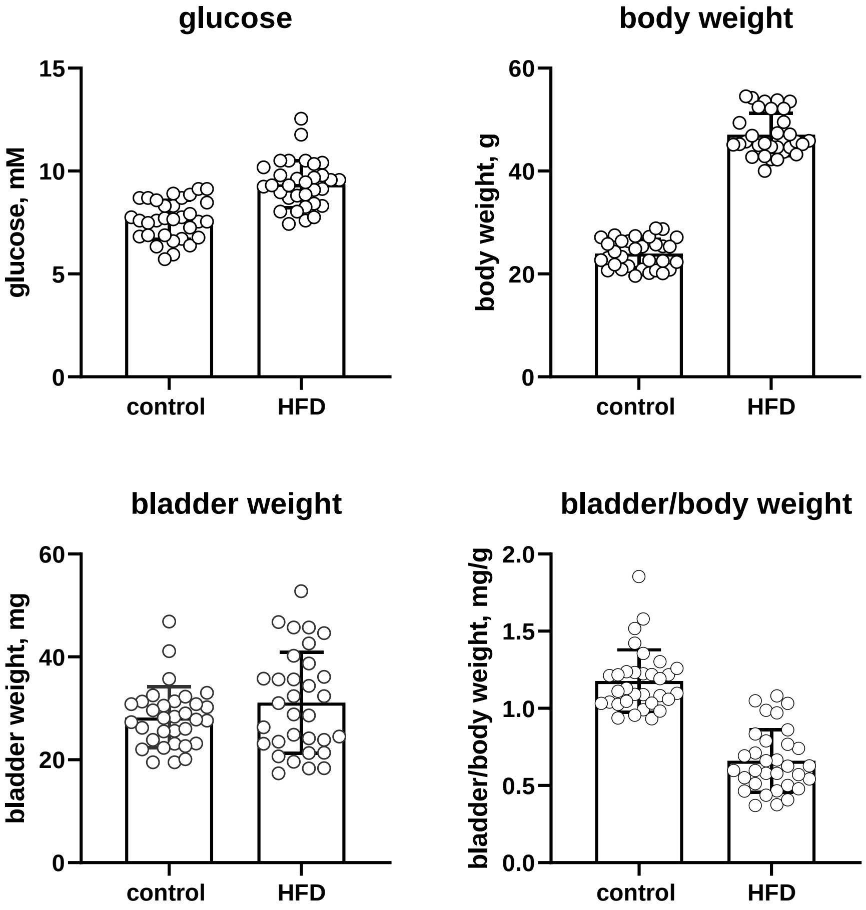

Supplemental Figure 26: Correlation of blood glucose with bladder and bladder/body weight in mice (C57BL/6N) on a control or HFD. Each data point represents one animal; the line represents the calculated regression line with its 95% CI. Descriptive p-values were 0.5655 and 0.1214, respectively.

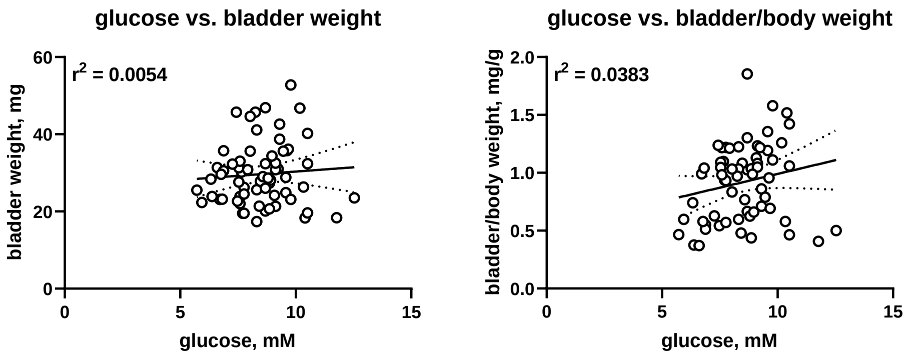


#### HFD mice + semaglutide treatment (Hoechst)

The study had been approved by the Ethics Committee of the State Ministry of Agriculture, Nutrition and Forestry, State of Hessen, Germany (the V54 - 19 c 20/15-FH/Anz. 1024). It was a 3-armed study: Male C67BL/6N mice (approximately 18-20 g body weight and 4-6 weeks of age) were obtained from Charles River (Sulzfeld, Germany) and placed on a control or an HFD for 18 weeks (EF adjusted fat diet, 25% fat content; ssniff Spezialdiäten, GmbH, Soest, Germany); thereafter, some of the mice with HFD received semaglutide (9) at a dose of 10 nmol/kg, every second day for another 36 days, i.e. approximately 29 weeks old at time of euthanization. Allocation to the three groups was based on randomization according to pre-treatment body weight. At study end, they were sacrificed by terminal bleeding from Vena cava caudalis under deep isoflurane anesthesia.

Supplemental Figure 27: Blood glucose, body weight, bladder weight and bladder/body weight in mice (C57BL/6N) on a control, HFD and HFD + semaglutide. Each data point represents one animal, bars and error bars represent means ± SD.

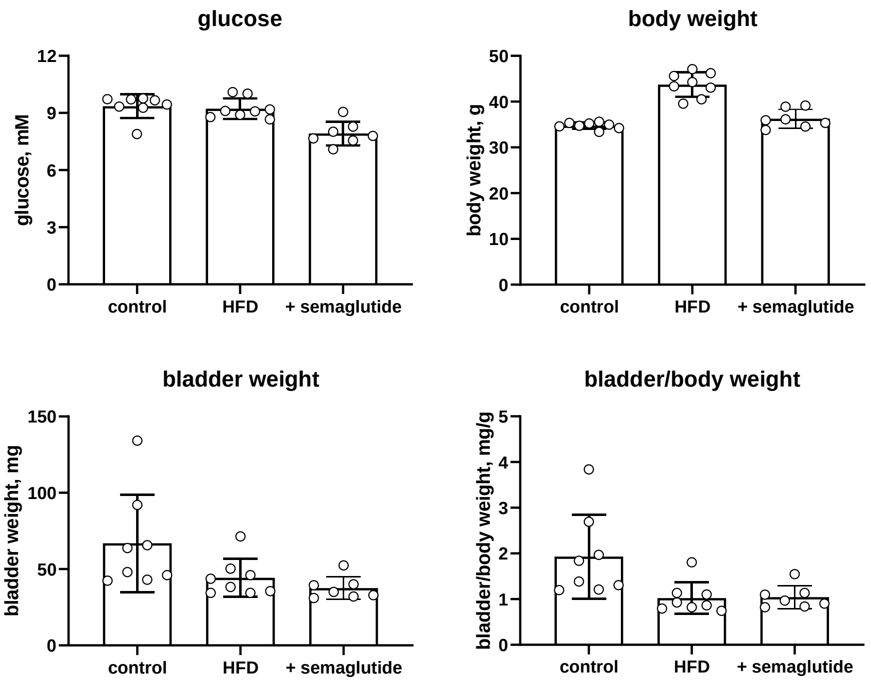

Supplemental Figure 28: Correlation of blood glucose with bladder and bladder/body weight in mice (C57BL/6JN) on a control or HFD. Each data point represents one animal; the line represents the calculated regression line with its 95% CI. Descriptive p-values were 0.1947, 0.2945, 0.2912 and 0.1322, respectively.

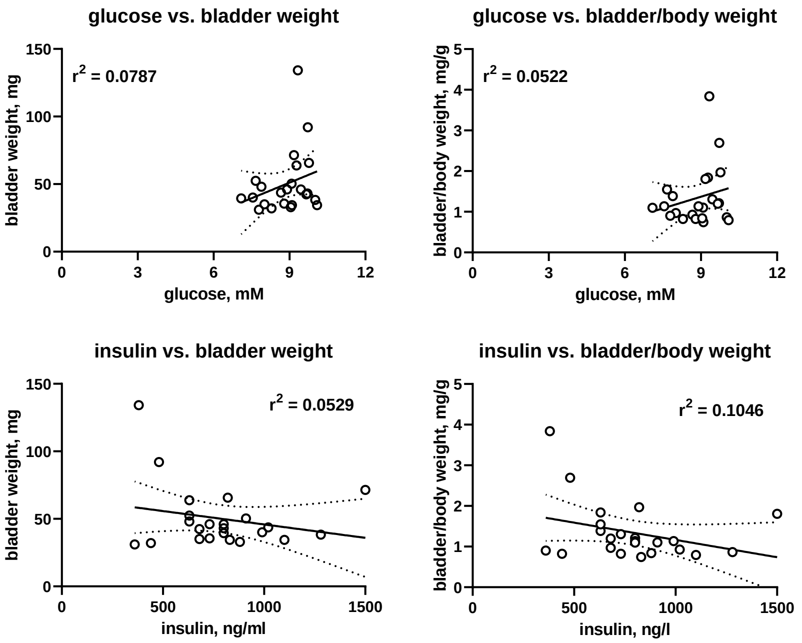


#### HFD mice (Mainz)

The study was approved by the responsible regulatory authority (Landesuntersuchungsamt Rheinland-Pfalz; 23 177-07/G 17-1-020). Male C57BL/6J mice were from Janvier Labs (Le Genest-Saint-Isle, France). Mice were fed ad libitum either with normal control or HFD for 21 weeks, beginning at the age of 6 weeks. The HFD (E15744-34, corresponding to Research Diets D12451) was obtained from ssniff Spezialdiäten GmbH (Soest, Germany) and was a defined, lard-based diet with 45% energy from fat, 35% from carbohydrates and 20% from protein. Twice a week food was exchanged, and food consumption was assessed. Mice were housed in cages with three to five mice.

Supplemental Figure 29: Fasting blood glucose, body weight, bladder weight and bladder/body weight in mice (C57BL/6J) on a control or HFD. Each data point represents one animal, bars and error bars represent means ± SD.

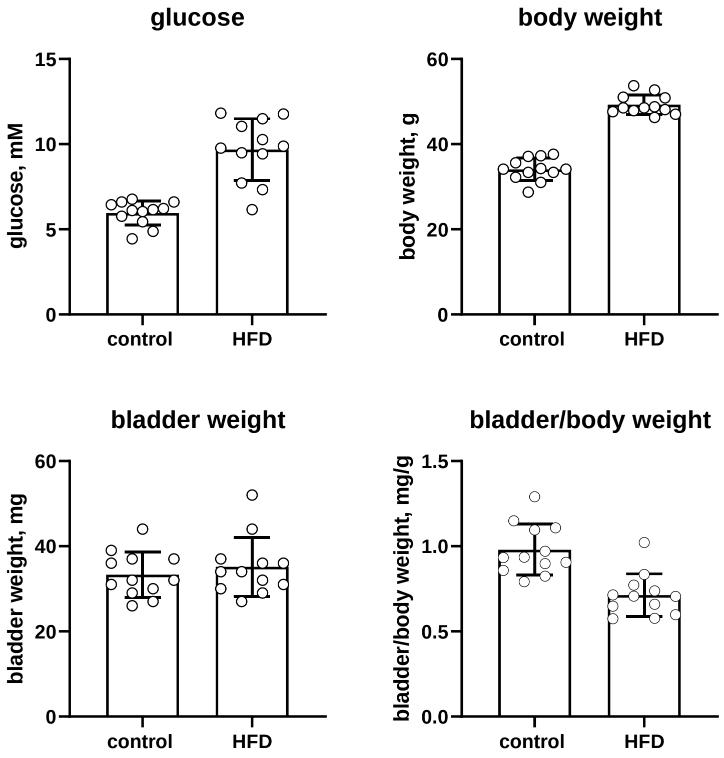

Supplemental Figure 30: Correlation of blood glucose and insulin with bladder and bladder/body weight in control (C57BL/6J) and HFD mice. Each data point represents one animal; the line represents the calculated regression line with its 95% CI. Note that the correlation between glucose level and bladder/body weight is primarily driven by the body weight gain on an HFD. Descriptive p-values were 0.4783, 0.019, 0.2459 and 0.0470, respectively.

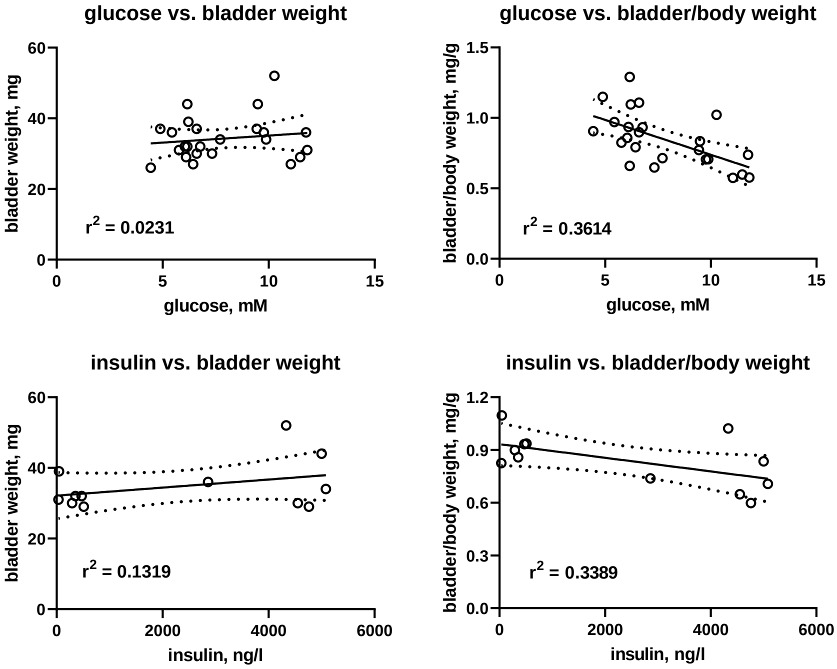
